# The telomere-to-telomere, gapless, phased diploid genome and methylome of the green alga *Scenedesmus obliquus* UTEX 3031 reveals significant heterozygosity and functional separation of the haplotypes

**DOI:** 10.1101/2022.11.30.518549

**Authors:** Thomas C. Biondi, Colin P.S. Kruse, Samuel I. Koehler, Taehyung Kwon, Wyatt Eng, Yuliya Kunde, Cheryl D. Gleasner, Kayley T. You Mak, Juergen Polle, Blake T. Hovde, Erik R. Hanschen, Shawn R. Starkenburg

## Abstract

Recent advances in sequencing technologies have improved contiguity of de novo genome assemblies. Nevertheless, the genomes of all eukaryotic organisms which are polyploid remain unfinished, limiting understanding of genetic and structural variation in diploid or polyploid organisms. Herein, we report the methodology and analysis of a 100% complete, gapless, phased, telomere-to-telomere diploid genome assembly of the eukaryote, *Scenedesmus obliquus* UTEX 3031 (DOE0152Z). Analysis of the fully assembled and resolved haplotypes revealed significant genomic rearrangements. Inter-haplotype heterogeneity was significant on most chromosomes yet one chromosome pair (Chromosome 15) was found to contain nearly no heterozygosity. Analysis of the 5mC methylation patterns revealed divergence in active gene content across haplotypes. Assembly of fully resolved chromosome pairs enabled complete resolution of genomic rearrangements and heterogeneity of haplotypes, the genomic basis of trait gain/loss, and evolutionary divergence across chromosome pairs. Further, when combined with 5mC methylation patterns, the assembly provides critical annotation information for genetic engineering approaches to achieve full knock-outs in allelic pairs.

## Introduction

Recent advances in sequencing technologies have increased sequence read lengths to tens to hundreds of kilobases, resulting in improved contiguity of de novo genome assemblies. When combined with physical maps (Bionano) and proximity ligation techniques (Hi-C), assembled genome fragments can be joined into chromosome scale scaffolds. Utilizing these and other techniques, Nurk, et al. successfully constructed the first gapless, telomere-to-telomere assembly for all chromosomes (sans the Y) of the human genome, revealing approximately 200 Mbp of novel sequence containing over 1900 novel gene predictions (Nurk et al., 2022). Additionally, Strand-seq has also been developed to sort and order contigs according to their haplotype. When combined with long-read sequencing data, Strand-seq has been used to assemble fully phased diploid genomes (Porubsky et al., 2021), further improving genome contiguity and completeness.

Despite these recent improvements, the genomes of eukaryotic organisms which are diploid or polyploid remain unfinished (Hanschen and Starkenburg, 2020).Complete assembly and phasing of each haplotype has yet to be achieved due to the complexity of long tandem repeats, collapsing of regions and phase switching between haplotypes that have low levels of heterogeneity, limiting our understanding of genetic and structural variation in polyploid organisms.

*Scenedesmus obliquus*, a single-celled, fresh-water photosynthetic eukaryote green algae, has been a model organism for fundamental photosynthesis research and was instrumental in the discovery of the Calvin-Benson-Bassham Cycle (Rieke and Gaffron, 1943; Benson and Calvin, 1948; Calvin and Benson, 1948; Lynch and Calvin, 1952; Buchanan et al., 1953). Furthermore, this species and other *Scenedesmus* are currently being used in biotechnology applications as sources of renewable fuels, wastewater treatment, and as a chassis for the biomanufacturing of commodity chemicals (Msanne et al., 2020). Unfortunately, the field lacks detailed knowledge of the *Scenedesmus* sexual cycle and its genetic and functional diversity, limiting the potential for genetic engineering, breeding and strain hybridization to further improve and identify industrially relevant traits. Additionally, given the high level of genetic diversity across algae (Hanschen and Starkenburg, 2020), it is often difficult to build structural gene models across different genera, which results in incomplete annotation, impeding the construction of accurate *in silico* metabolic models that propel strain improvement. To this end, a PacBio-based draft genome assembly of *S. obliquus* UTEX 3031 (Starkenburg et al., 2017) was constructed, indicating that this microalgae was a natural diploid. Sequencing of additional genomes from the genus *Scenedesmus* have revealed that this genus contains both haploid and diploid members (Polle and Starkenburg, unpublished results), strengthening the hypothesis that this genus is capable of sexual reproduction.

To enable breeding, trait mapping, and to acquire strain performance-information that cannot be garnered from incomplete genomes, we initiated a genome finishing project to fully assemble the genome of *S. obliquus* UTEX 3031. Herein, we present the methods and initial analysis of the complete, gapless, phased, telomere-to-telomere diploid genome assembly of the photosynthetic eukaryotic algae, *S. obliquus* UTEX 3031. To our knowledge, this is the first 100% completely phased diploid assembly for any eukaryote. Comparative analysis of the resolved chromosome pairs revealed large genomic rearrangement between haplotypes, suggesting the haplotypes are actively evolving specialized functions. Furthermore, additional evidence supporting sexual recombination was found, candidates for sex chromosomes were identified, and CpG methylation information was used to inform differential gene activation between haplotypes. Additional analysis of these and other fully phased and complete polyploid genomes will contribute to our understanding of genetic diversity, structural variation, and enable the ability to decipher the full range of genome variation/functional diversity that have evolved in polyploid organisms.

## Results

### Genome finishing and gap closure

The draft genome assembly of *S. obliquus* UTEX 3031 (Starkenburg et al., 2017) was improved and scaffolded using a combination of PacBio HiFi, Oxford Nanopore, and Illumina Hi-C reads. Each read type was incorporated serially to assess the impact of each technology on genome contiguity (Table 1). HiFi reads alone greatly improved the contiguity of the assembly, increasing the N50 from 155 Kbp to 4.069 Mbp, reducing the number of contigs from 2,705 to 204, and increasing the maximum contig size to 11.144 Mbp; the full telomere-to-telomere assembly of the largest chromosome (chromosome 1A.) Of note, Bionano maps were also constructed and evaluated. However, due to the low density of restriction sites, minimal improvement in contiguity/scaffolding were observed (data not shown). In this assembly, the Hi-C data was critical to orient the contigs into chromosome-level, phased scaffolds. In addition, Hi-C sequencing reads helped identify ‘false’ contigs, or contigs that did not seem to scaffold into any particular chromosome due to contamination or small repeat sequences that could not be confidently placed in the genome. Lastly, gap-closing with Oxford Nanopore reads reduced the scaffold count to 34 – representing telomere-to-telomere scaffolds for each phased chromosome.

**Table 1.** *Scenedesmus obliquus* UTEX 3031 Genome Assembly Improvements.

| Assembly | Initial Draft (2017) | Draft 2 | Draft 3 | Final Assembly | Difference from Initial Draft |
| --- | --- | --- | --- | --- | --- |
| Sequencing Data Types | PacBio CLR | PacBio HiFi | PacBio HiFi + Illumina Hi-C | PacBio HiFi + Illumina Hi-C + Oxford Nanopore |  |
| # of Contigs | 2,705 | 204 | 68 | 34 | -98.7% |
| Contig N50 | 155,544 bp | 4,069,430 bp | 6,027,826 bp | 6,027,792 bp | +3,775% |
| Size | 207,967,116 bp | 207,501,956 bp | 202,415,811 bp | 205,222,550 bp<sup>+</sup> | -1.32% |
| Largest contig | 2,334,183 bp | 11,144,349 bp | 11,144,349 bp | 11,144,322 bp | +377.4% |
| Complete BUSCOs | 80% | N.D. | N.D. | 97.4% | +21.8% |

<sup>+</sup> Includes mitochondrial and chloroplast genomes

After combining the sequencing information from all technologies, approximately 34 gaps remained in the genome. Four false duplications and five mis-assemblies were fixed by performing a local reassembly with hifiasm. To fill the 22 gaps caused by long tandem repeats (LTRs), we followed the approach by Nurk *et. al. (Nurk et al., 2022)*, using both CCS reads and ONT reads. The three gaps caused by (TG)_n_ dinucleotide repeats were fixed with PacBio HiFi reads and corrected Nanopore reads spanning these gaps. As a final quality check, we analyzed and compared the HiFi read coverage across the entire genome before and after gap closure to confirm uniform coverage across each chromosome (Supplemental Figure S1). Collectively, approximately 3 Mbp of new sequence was added to the assembly to correct long-tandem repeats that were incorrectly collapsed.

### Genome Overview

Post gap closure, the complete *S. obliquus* UTEX 3031 genome consists of 205,222,550 bp contained in 34 chromosomes (2n=34), a mitochondrial, and a chloroplast genome, resulting in a Merqury (Rhie et al., 2020) QV score of 54.6. Using DNA staining and flow cytometry, we predicted the genome size of UTEX 3031 single cells, finding the sequenced diploid genome size of 205 Mbp is consistent with measured DNA content from flow cytometry (Supplemental Figure S2). BUSCO analysis further demonstrated that each haplotype of UTEX 3031 (denoted as A and B) contain nearly complete BUSCO gene sets (97.4%, Supplemental Figure S3) providing high confidence that all genetic content has been captured in the assembly. To quantify the degree of heterozygosity, the average nucleotide identity (ANI) between chromosome haplotypes was calculated and ranged from 92.3% to 100%, with the vast majority of pairs falling between 94.5% and 96.5% (Figure 1). However, there are a few outliers of note. Chromosome 15 has an ANI of 100%, indicating that the two haplotypes are near exact duplicates of one another. In addition, chromosome 16 has the lowest ANI (92.3%) of any chromosomal pair in the genome.

**Figure 1.**
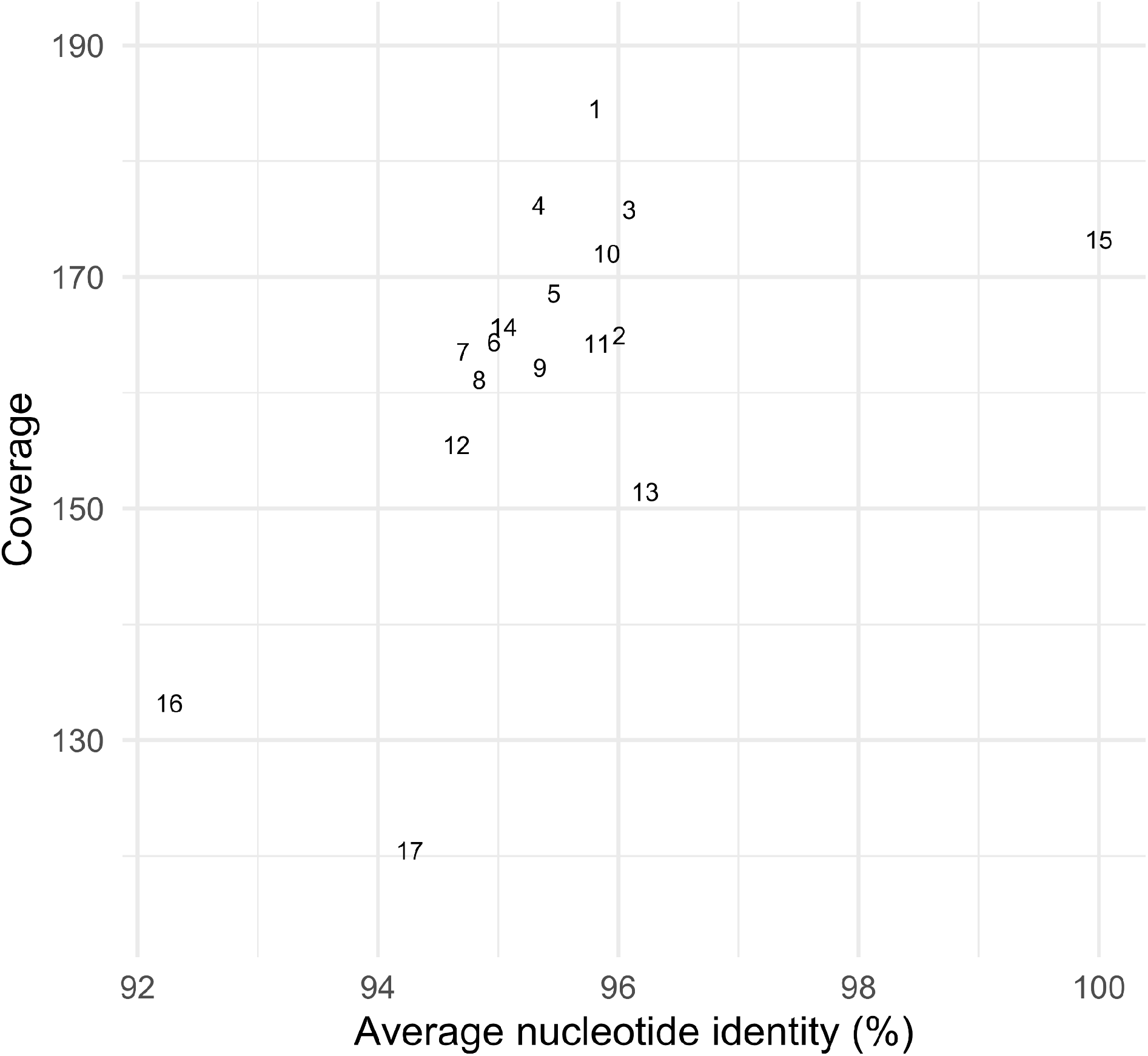
Average Nucleotide Identity (ANI) vs read coverage for each chromosome. PacBio HiFi read coverage for reads only aligned to haplotype A versus the ANI between the A and B haplotypes for each chromosome pair.

To gain a better understanding of the high degree of synteny observed between the chromosome 15 haplotypes, we measured the average PacBio HiFi read coverage for each chromosome in the genome. Collectively, read coverage ranges from 118x coverage to 182x coverage, with chromosome 15A having an average coverage of 149x. These data suggest that two copies of chromosome 15 are maintained vs. a loss of one haplotype resulting from an aneuploidy event. Furthermore, through comparative analysis with other sequenced *Scenedesmus* strains, homologous chromosomes were identified in the closely related *Scenedesmus obliquus* DOE0013 genome (scaffold 5) which is near chromosome length with few rearrangements compared to chromosome 15 in UTEX 3031. Similarly, homologous scaffolds for chromosome 15 were found in the Scenedesmus obliquus EN004 genome assembly (Supplemental Figure S4). However, we were not able to identify homologous chromosomes in the closest sequenced relatives of *Scenedesmus* (*Raphidocelis subcapitata* and *Monoraphidium neglectum* (Suzuki et al., 2018)), nor the more distantly related *Chromochloris zofingiensis*. Based on current information, the data suggests that chromosome 15 in UTEX 3031 consists of two near-identical haplotypes. The mechanism for this loss of chromosomal variation is not known and warrants further investigation.

With respect to gene content, the genome encodes 29,226 characterized and putative genes, 47% (13,839) of which could be assigned some level of functional annotation (Table 2). To determine the contribution of individual chromosomes to total coding sequences, each chromosome’s gene quantity and length were calculated (Supplemental Figure S5), indicating that the larger chromosomes have a higher coding density than the smaller chromosomes. Collectively, the ‘A’ haplotypes, defined as the larger of each chromosome pair across all chromosomes, contain 250 more genes than the ‘B’ haplotypes.

**Table 2.**
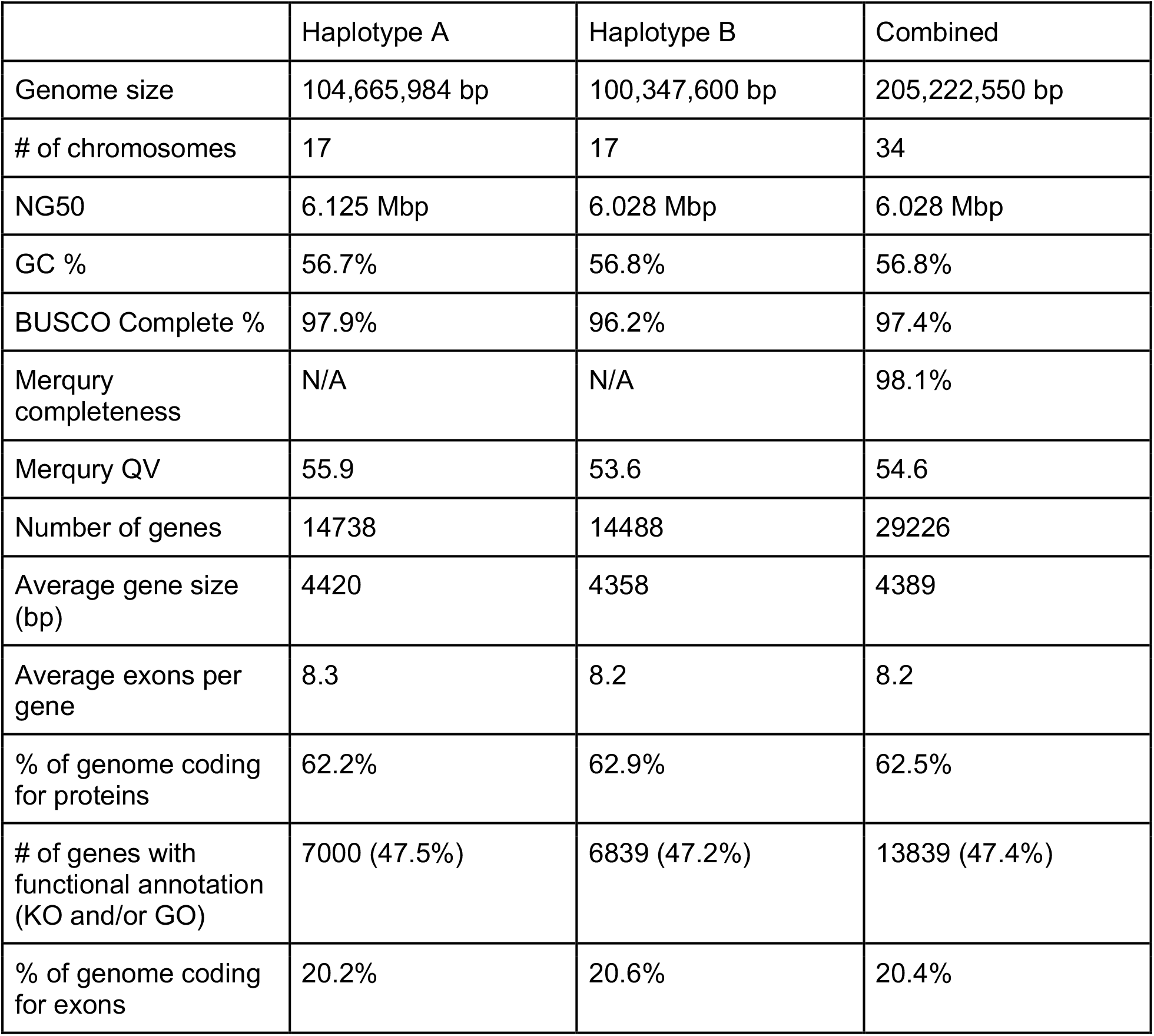
Genome statistics of *Scenedesmus obliquus* UTEX 3031

Large deletions in two chromosomes were also observed over the course of the project (DNA was extracted from culture samples for Illumina sequencing approximately 3 years prior to DNA extracted to generate the PacBio HiFi reads). When inspecting the mapping coverage of Illumina and PacBio HiFi reads across the genome, anomalies were detected in two specific regions (Supplemental Figure S6). In each of these two regions, the sequencing coverage was doubled in the region of interest compared to the rest of the chromosome only in the Illumina sequencing data. Close inspection revealed that in both cases, a large chromosomal deletion had occurred in one of the haplotypes in the intervening time between DNA sequencing experiments. These alterations in genome topology were confirmed/verified by PCR (Supplemental Figure S7).

### Synteny and Genomic Rearrangements

A fully phased, diploid genome provides a unique ability to understand the structural rearrangements between haplotypes in diploid species. In the *S. obliquus* UTEX 3031 genome, there are three major indels (>500 Kbp on chromosomes 2, 9, and 13) between haplotypes (Figure 2). To evaluate the differential function induced by these changes, gene content was analyzed (Supplemental Figure S8). There are two large-scale rearrangements on Chromosome 2A, one an expanded ribosomal RNA array (Figure 2, blue duplication). The other (Figure 2, indel near 7 Mbp) region in Chromosome 2A contained a significant enrichment for N,N-dimethylaniline monooxygenase and flavin adenine dinucleotide activities and represented one third of the annotated genomic capacity for N,N-dimethylaniline monooxygenase activity (Supplemental Figure S8). The region in Chromosome 9A contained a significant enrichment for structural constituents of chromatin and DNA binding activities and contained the only two genes annotated with phosphatidylserine decarboxylase (PSD) capacities. The region in Chromosome 13A did not reflect any obvious consequence with the only significantly enriched functionality being the broad protein binding term (Supplemental Figure S8).

**Figure 2.**
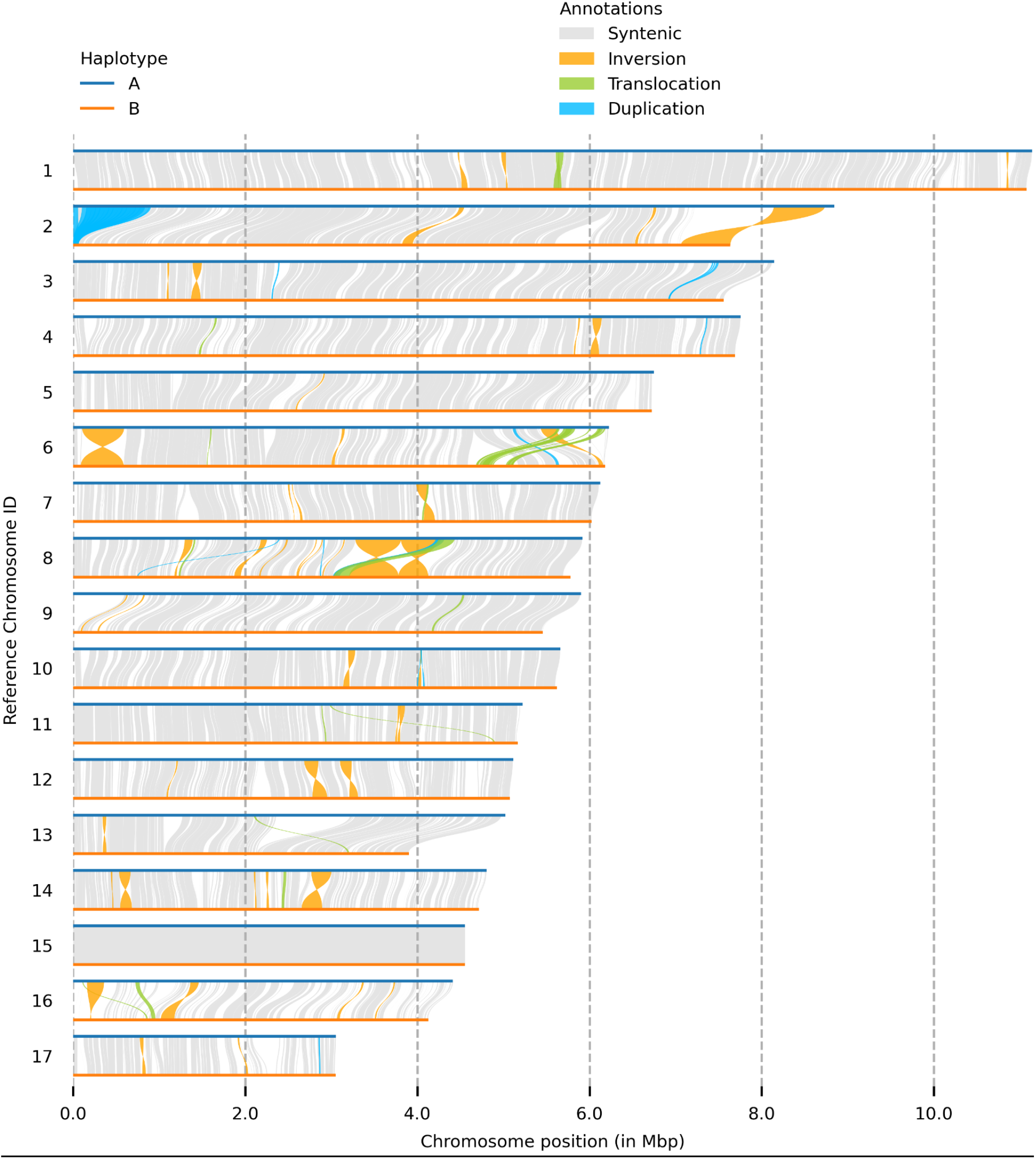
Chromosome scale genomic rearrangements and synteny between haplotypes.

The proportion of BUSCO genes within the indel regions ranged from 6% of genes in chromosome 2A to 13% in 9A, suggesting these regions represent a loss of necessary functions from one chromosome pair. To identify the potential effects of this heterozygosity, the methylation state of the retained BUSCO alleles in each indel were evaluated (see CpG methylation methods and results below). Across all three deletions, every BUSCO gene was found to have a hypomethylated promoter region, indicating active expression and implying that the extant allele is being expressed as necessary to retain necessary functions in the diploid UTEX 3031. In keeping with this trend, both of the promoter regions of the aforementioned PSD genes that are exclusive to the indel region of 9A indicate active expression (hypomethylation).

To understand smaller genomic rearrangements (between 500 bp and 500 Kbp), a detailed comparative analysis was performed to identify all genomic rearrangements (GRs), syntenic regions between each haplotype for all chromosomes, and the unaligned regions that do not belong to any of GRs or syntenic regions (Figure 3A). GR regions were classified into two groups based on the reference sequence: inter-chromosome and inter-haplotype. Inter-chromosome GRs refer to rearrangements from one chromosome to an unpaired chromosome (e.g. chromosome 1A to chromosome 5B), whereas inter-haplotype GRs refer to rearrangements within a chromosome pair (e.g. chromosome 1A to 1B). GRs, regardless of location, were also further categorized into five classes: Inversion (INV), Translocation (TRANS), Inverted translocation (INVTR), Duplication (DUP), and Inverted duplication (INVDP).

**Figure 3.**
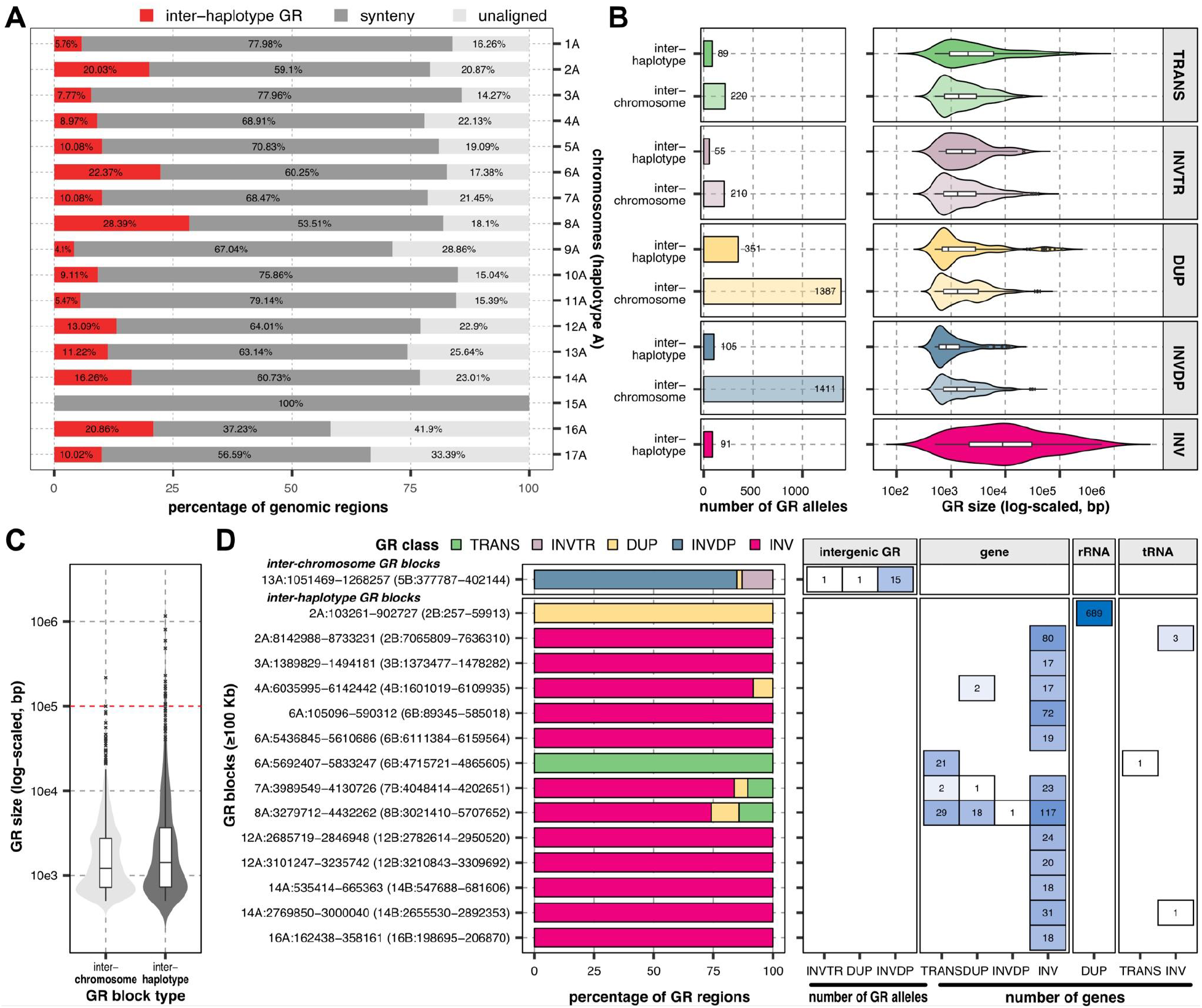
Summary of genomic rearrangements. (**A**) composition of synteny and inter-haplotype genomic arrangements. (**B**) numbers and sizes of genomic rearrangement alleles depicted by class. (**C**) Sizes of genomic rearrangement blocks. Red dashed line indicates the size cutoff for long genomic rearrangement blocks. (**D**) Quantification of genomic rearrangement allele regions (left panel) and composition of genes (right panel) in genomic rearrangement blocks larger than 100 Kbp. Colored bar (left panel) indicates genomic rearrangement class. Genomic rearrangement block name indicates its query region in parentheses. TRANS indicates translocation. INVTR represents inverted translocation. DUP indicates duplication. INVDP indicates inverted duplication. INV indicates inversion.

On average, only 68.0% of all chromosome pairs were syntenic (at >90% identity cutoff), ranging from 37.2% between haplotypes of chromosome 16 to 100% of chromosome 15 (**Figure 3A)**. On average, 11.8% of a given haplotype has been re-arranged between haplotype A and B (inter-haplotype GR), ranging from 0% in chromosome 15 to 28.4% in chromosome 8A. A significant correlation was not measured between proportions of syntenic regions and chromosome sizes (Spearman’s rho=0.27, *p*=0.29). Broadly, a total of 3,920 GRs (500 bp or more in length) were detected, comprising 3,229 inter-chromosome GRs and 691 inter-haplotype GRs (**Figure 3B**), the majority of which were duplications (DUP=1738 and INVDP=1516). The sizes of GR alleles ranged from 3083.81 bp (INVDP) to 98024.67 bp (INV), indicating large variability in GR allele sizes. Inter-chromosome INVs were not detected due to the sequence context between different chromosomes (**Figure 3B**). However, the size of the inversions (INVs) between two haplotypes of a chromosome have the highest median size (3,672 bp) among all GR classes.

To characterize the largest rearrangements across the genome, we further investigated consecutive spans of GRs, termed ‘GR blocks’ (**Figure 3C)**. In total, we found 3,018 GR blocks, comprising 2,534 inter-chromosome blocks and 484 inter-haplotype blocks. On average, inter-chromosome GR blocks were smaller than inter-haplotype GR blocks with median sizes of 1203 Kbp and 1421 Kbp (Wilcoxon Rank Sum test *p* = 0.0021), respectively (**Figure 3C**).

The largest GR blocks (n=15), spanning 100 Kbp or more, are depicted in **Figure 3D**. We found a single inter-chromosomal GR block on chromosome 13A (13A:1051469-1268257) which primarily consists of an inverted duplication from chromosome 5B (**Figure 3D**). Among 14 inter-haplotype GR blocks, 12 of them were enriched with large inversions (**Supplemental Figure S9**). One inter-haplotype GR block on chromosome 2 (2A:103261-902727) is the result of variation in the copy number of the rRNA genes: a ∼8 Kbp region in chromosome 2B that is repeated approximately 105 times on chromosome 2A. (**Figure 3D**).

To further examine the genomic rearrangements and features of interest, an integrated analysis of methylation, gene content, translocations, and repeat regions was performed (Figure 5). Examination of gene translocation events show chromosomes 10 and 16 demonstrate many gene translocations on both haplotypes. Only chromosome pair 15 does not have detectable translocations, consistent with 100% identity of chromosome 15a and 15b (Figure 1). When comparing rates of intra-chromosomal rearrangements between homologous chromosome pairs, chromosomes 2, 4, 13, and 17 have the most divergent propensities for translocations. Chromosomes 4A, 13A, and 17B demonstrate greater likelihood of translocation events compared to their chromosome pair (Supplemental file 1, Figure 5).

### CpG methylation & Gene Activation Analysis

CpG methylation, or transformation of cytosine into 5-methylcytosine in a CG motif context, has been shown to play a role in regulating gene expression (Razin, 1998). A gene can be transcribed if the promoter is unmethylated, but can be silenced or repressed by methylating the promoter, making it inaccessible to transcription factors (Deaton and Bird, 2011). By extracting the CpG methylation information from PacBio HiFi read files and overlaying the CpG methylation probabilities with the structural gene models, we observed that the cytosine bases in the coding regions of the gene models are highly methylated while the regions immediately upstream of the annotated start site of a gene are characterized by hypomethylation (Figure 4). This pattern of reduced methylation in the putative gene promoter regions is present for 89.3% of the annotated genes while 11.7% (3,408) of genes lack hypomethylation signatures upstream of transcription start sites (TSS). A median distance of 677 bp was observed between a given TSS and its nearest putative promoter, while the median promoter region was calculated to be 803 bp wide. Due to the lack of heterozygosity, methylation probabilities could not be calculated between chromosomes 15A and 15B.

**Figure 4.**
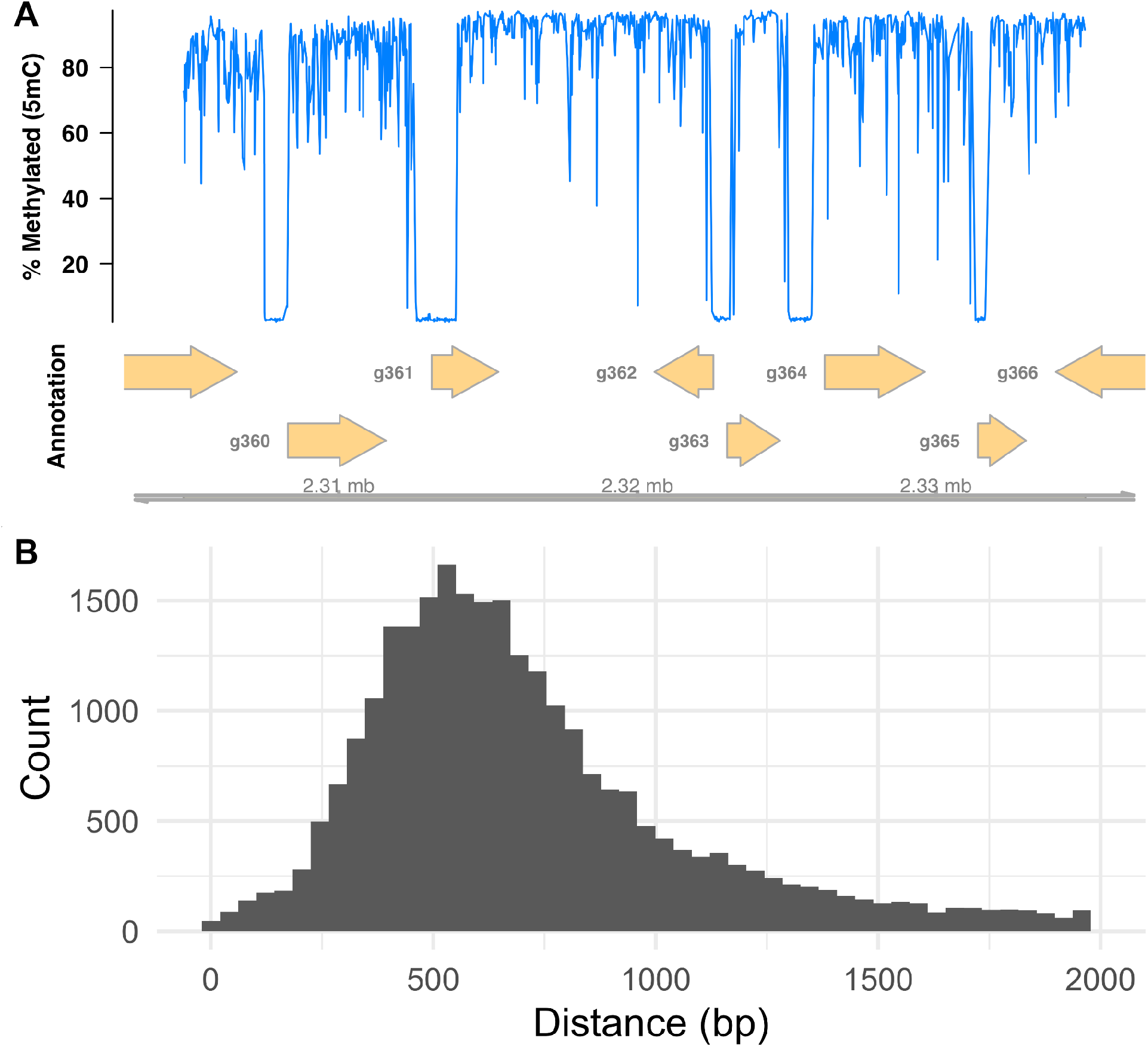
A) Example of the relationship between regions before the start of genes and hypomethylated regions. B) Histogram of distances from gene start sites to putative promoter start sites.

Interestingly, for the genes containing methylated promoter regions, three annotation terms were found to be enriched, including genes associated with regulation of viral transcription (*p* = 0.0029), double-strand break repair via homologous recombination (*p* = 0.0029) and DNA integration (*p* = 0.0136) (Supplemental Figure S10). Among the methylated genes associated with regulation of viral transcription, a homolog of magnesium chelatase subunit H 0 (bchH0) was hypermethylated in a manner consistent with the silencing of bchH0 to facilitate viral attack in higher plants (Hiriart et al., 2003). To further understand the regulatory nature of these methylation events and the degree of functional divergence between the haplotypes, we analyzed all gene paralogs that shared >90% shared sequence similarity (6,648/29,226 (22.8%) of genes or 3,324 gene pairs). Of these 3,324 pairs, the promoter regions of (87.9%) both alleles were not methylated, 180 (5.4%) were methylated in both genes, and 223 (6.7%) were observed to be differentially methylated (the promoter region of one allele was methylated while the 2nd allele was unmethylated) (Supplemental Figure S11). The function of the alleles containing differential methylation patterns were found to be overrepresented with genes associated with DNA integration (*p* = 0.0006), while gene pairs where both alleles were methylated had no significantly enriched terms.

A variety of methylation states were found across all chromosomes, but Chromosome 2 showed several unique and biologically interesting features. The expanded ribosomal RNA array (Figure 2) contained low levels of methylation which suggest that most copies of the rRNA genes are actively transcribed (Figure 5,*). Furthermore, the opposite end of chromosome 2B was observed to contain a region with noticeably higher gene density/hypermethylation relative to all other regions of the genome (Figure 5,**). This region was flanked by two repeat regions on haplotype 2B. Analysis of the 50 genes contained in this region identified 31 proteins of unknown function, the largest grouping of genes. However, the remaining genes shared the highest degree of homology with viral (12 genes) or bacterial associated genes (4 genes) with only three genes annotated as eukaryotic in origin. The region is simultaneously one of the most gene dense regions in the genome which may indicate a higher propensity for gene insertion events or that the region is the site of a large historical gene insertion.

**Figure 5.**
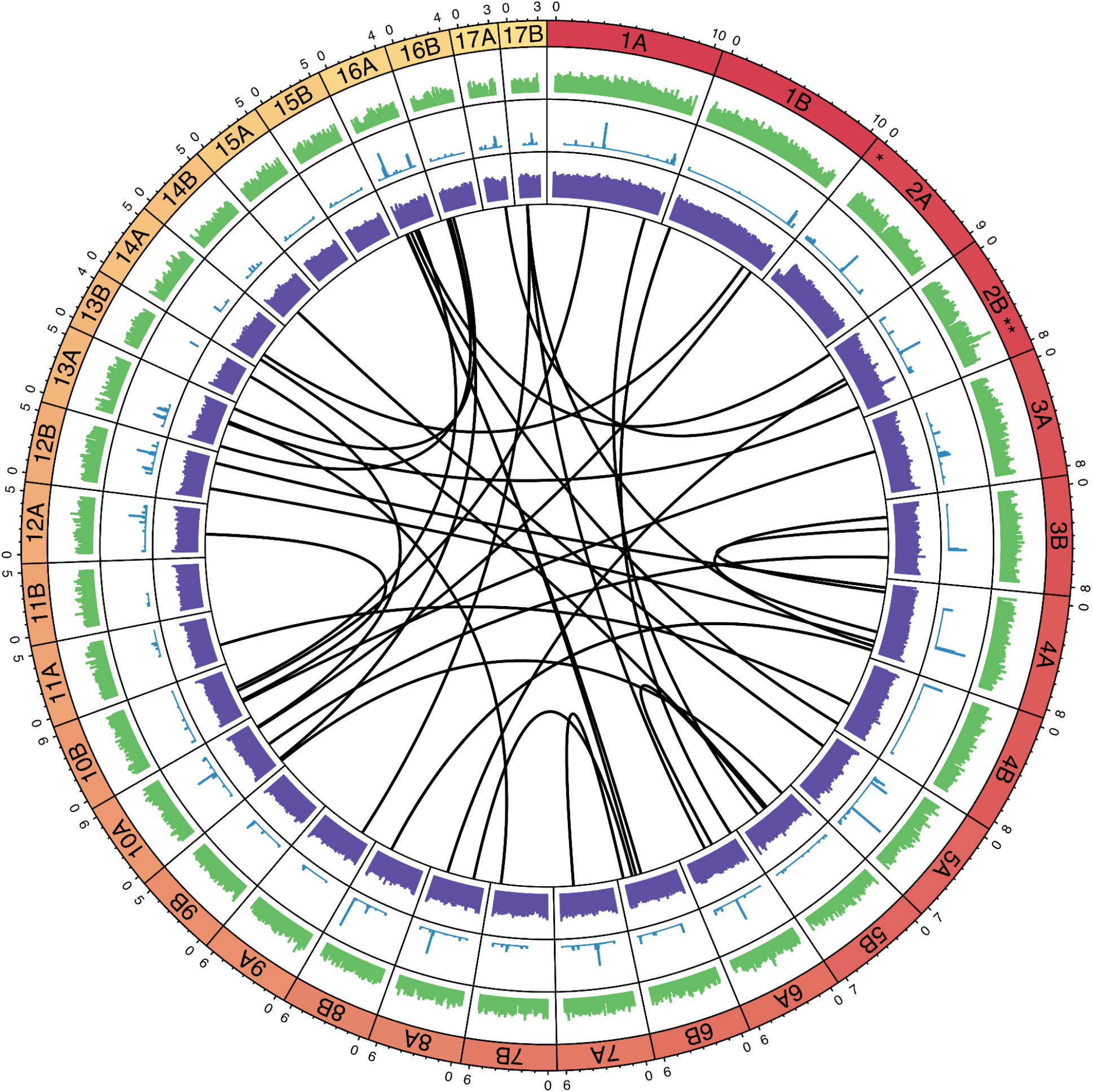
Haplotype Resolved Chromosome Features. Chromosomes are indicated by the Red-Yellow gradient, with outer numbers indicating size (Mbp). The gene density (green), repeat regions (blue), methylation probabilities (purple) and putative gene translocations (black lines) are depicted. Symbols ‘*’ and ‘**’ in chromosome bands identify hypomethylated rRNA duplication region and gene-dense, hypermethylated region with high viral and bacterial gene content respectively.

### Sexual recombination and sex chromosome analysis

Given the interest in leveraging breeding strategies to improve traits in algal lineages, and that several reports indicate that sexual reproduction occurs in *Scenedesmus* (Francis R. Trainor, 1963; Trainor and Burg, 1965; Cain and Trainor, 1976; Hindák and Trainor, 1995; Trainor, 1996; Cepák et al., 2006) and in other Chlorophyceae green algae (Ferris et al., 2010), we completed several analyses to search for evidence of sexual recombination and the presence of sex chromosomes in UTEX 3031. First, we looked for evidence of sexual reproduction and recombination amongst *Scenedesmus* strains by examining the relationships of genes amongst *Scenedesmus* strains. However, the comparative analysis of the phylogenetic signal among multiple *Scenedesmus* strains using gene coding regions or exons indicate that the gene identities are nearly identical and therefore cannot be used to determine relatedness of UTEX 3031 to other strains (data not shown). Thus, we next compared predicted introns against other *Scenedesmus* strains, finding numerous genomic regions where introns are closest to either a single strain or to numerous strains (Figure 6). Some regions of introns were observed to be most closely related to EN004 (which had the most introns with highest similarity to any given strain) while other regions of introns are more closely related the remaining strains, which is indicative of recombination between strains during sexual reproduction. Third, we attempted to identify sex-restricted/sex-determining chromosomes by searching for the location of genes containing RWP-RK domains (Supplemental Figure S12), which have been shown to be associated with Chlorophycean sex chromosomes (Umen, 2020). Genes annotated with RWP-RK domains (PF02042 or IPR003035) were found on chromosomes 4, 6, and 12. Fourth, we analyzed the GC content of introns on all chromosomes as the GC content of introns on incipient sex chromosomes can be elevated due to reduced recombination and GC biased gene conversion (Charlesworth et al., 2020). Focusing on chromosomes 6 (due to its increased rearrangements), 15 (due to its increased homozygosity) and 16 (due to its increased heterozygosity) (Figure 2), we found that chromosome 6A has a noticeable decrease in GC content beginning at ∼6 Mbp (Supplemental Figure S13), which corresponds to the region of this chromosome with increased rearrangements (Figure 2). Lastly, we looked at chromosome level methylation, finding no large region of methylation which may indicate a transcriptionally repressed region during asexual growth and reproduction (when cells were harvested for DNA sequencing) (Figure 5).

**Figure 6.**
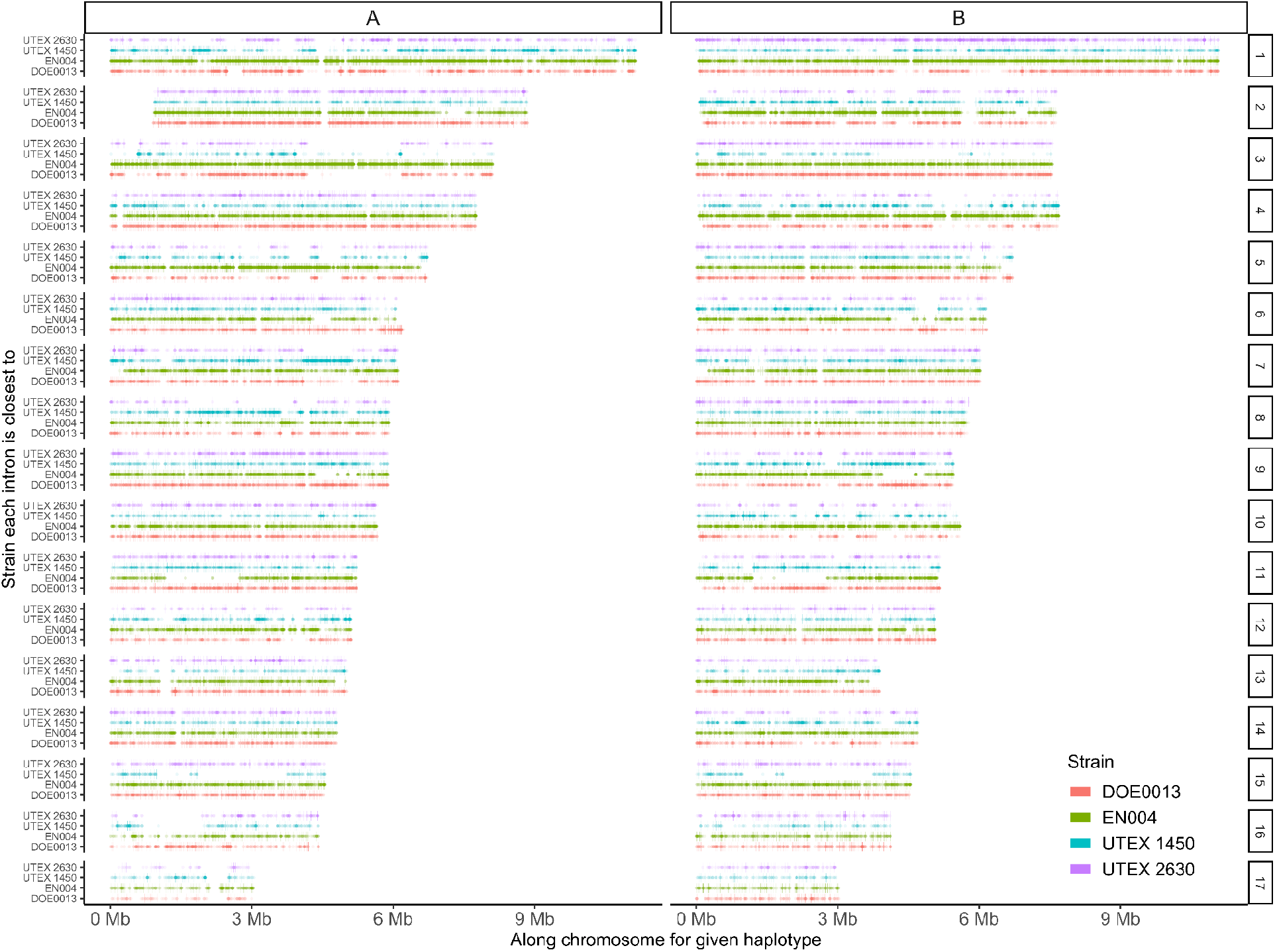
Intron analysis. Location of each intron and which other strain of *Scenedesmus* each intron is most closely related to (DOE0013, EN004, UTEX 1450, UTEX 2630). Various regions show clear similarity to only one strain (e.g. chromosome 3A at approximately 4-6 Mb shows close relationship to strain EN004 whereas chromosome 7A at approximately 4-5 Mb shows close relationship to strain UTEX 1450).

## Discussion

Here, we report the generation of the first telomere-to-telomere, gapless, haplotype-phased diploid genome assembly for an industrially relevant eukaryotic alga. While several complete assemblies have been generated for haploid organisms (Nozaki et al., 2007; Lemieux et al., 2019; Nurk et al., 2022; Giguere et al., 2022) prior to this study, fully phased polyploid genomes had yet to be assembled. Importantly, the availability of this fully phased gapless genome enabled novel discoveries that were not possible with the initial (yet still highly contiguous) draft assembly. First, full assembly of both haplotypes enabled the identification of multiple, significant large-scale genomic rearrangements between haplotypes. Specifically, we identified a large duplication in rDNA copy number between chromosomes 2A and 2B, a ∼1 Mb deletion in chromosome 2B, and a ∼1 Mb deletion in chromosome 13B. Second, measurement of the differential hypomethylated regions upstream of allelic gene pairs enabled the ability to quantify functional specialization/separation between haplotypes. Third, the full diploid assembly enabled the identification of temporal shifts in genome content on two chromosomes, critical information that can help monitor trait gain/loss during long term lab maintenance. Furthermore, we were also able to discover a complete loss of heterozygosity in *S. obliquus (*chromosome 15). A similar, unexplained event has been found in chromosome 19 of *Phaeodactylum tricornutum*, an unrelated, evolutionarily distinct marine diatom (Giguere et al., 2022). The significance and frequency of loss of heterozygous haplotypes across other algae remains to be determined. We anticipate that this is the first of many complete higher-ploidy genome assemblies to enable the full exploration and characterization of large-scale structural rearrangements and the evolution of functional diversity between haplotypes.

The observed connection and consistency between the putative promoter regions and their lack of 5mC methylation can be leveraged going forward to improve *de novo* structural annotation pipelines. The methylation data also revealed a unique region in chromosome 2B with evidence of potential de novo methylation to silence a variety of viral genes (Šenigl et al., 2012) which was further supported by an enrichment of genes related to viral DNA integration among the hypermethylated promoters. The region in chromosome 2B is of additional interest as one of the most gene dense regions in the genome. The combined features of this repeat-bounded region on chromosome 2B may indicate the region is maintained as highly available to foreign DNA integration, which warrants further investigation. These methylated regions begin to provide new insights into the mechanisms algae use to silence foreign DNA, as previously observed in *Chlamydomonas (Neupert et al., 2020)*.

Collectively, our analysis also supports the hypothesis that sexual reproduction is occurring in *Scenedesmus*; neutrally evolving introns in *S. obliquus* UTEX 3031 appear to show recombination along chromosomes (Figure 6). However, with the available complete genomic information from a single strain of *S. obliquus*, we are not currently able to conclusively identify any incipient sex chromosomes. Several chromosomes are candidates, due to increased rearrangements or increased heterozygosity, but the presence or absence of RWP-RK protein domains (Supplemental Figure S12) should not be considered necessary and sufficient evidence for or against any chromosome being a sex chromosome as *S. obliquus* may or may not use the same domain to regulate sexual reproduction as other green algae (Umen, 2020). Similarly, the GC content of predicted introns may not be affected by lack of recombination due to a relatively young sex-loci age (Ferris et al., 2010). Other chromosomes, such as chromosome 16 (with increased heterozygosity) may be sex-determining chromosomes or contain sex-determining regions, but with the available information, the sex-determining chromosome in *Scenedesmus obliquus* cannot be conclusively identified. Furthermore, it is unknown whether *Scenedesmus* is homothallic (a single genotype produces two different gametes usually capable of fusion) or heterothallic (two genotypes exist, each producing different gametes not capable of self-fusion), complicating the identification of sex-loci in *Scenedesmus*. Additional high quality genomes of *Scenedesmus* would greatly benefit the identification and understanding of sexual reproduction and sex loci in this important Genus.

As a practical note, the increasing availability of fully phased polyploid assemblies should prompt community discussion about database standards. For example, NCBI only allows diploid assemblies to be deposited as two separate accessions for a given organism. This structural limitation for polyploid assemblies in public databases requires additional procedures for use, which may complicate community accessibility and limit genome utilization.

## Methods

### Culturing and DNA Extraction

UTEX 3031 (previously known as DOE0152Z) was cultured in BG11 media (https://utex.org/products/bg-11-medium) at 25°C in a Percival incubator programmed with a 16hr/8hr diel cycle. Once the culture reached early stationary phase, the cells were harvested, centrifuged for 10 minutes at 3,500rpm/4°C and stored at -20 C prior to DNA extraction. Cells from the frozen algal pellet were thawed, washed with 3 mL of BG-11 buffer, centrifuged 10 minutes at 3,500rpm/4°C. The supernatant was decanted, and cells were resuspended in a resuspension buffer (200mM NaCl, 100mM EDTA, 10 mM TRIS, pH 7.2). An equal volume of 1% LMP agarose solution was added to the cell suspension and mixed gently by swirling the tube. The mix was pipetted into the plug molds and incubated at 4°C to solidify. Resulted agarose plugs were incubated in the protoplasting solution overnight at 37°C with gentle mixing to remove the cell wall. Next, the plugs were incubated overnight in the lysis buffer with Proteinase K. Lyzed plugs were washed three times with 1xTE buffer and digested overnight with beta-Agarase I to release the gDNA into the solution. gDNA was purified according to the High Salt:Phenol:Chloroform:IsoAmyl Alcohol protocol recommended by Pacific Biosciences. Purified gDNA was concentrated with AMPure PB beads and was used to prepare various PacBio SMRT bell libraries and for Nanopore sequencing.

### Estimation of Nuclear DNA Content

The ploidy level/DNA content of *Scenedesmus obliquus* UTEX 3031 was estimated by comparing to other algal cultures with known genome sizes; *Nannochloropsis oceanica* (P7C12 (an internal strain designation), CCAP 849/10, and CCMP1779), *Chlamydomonas reinhardtii* CC-124, and *Dunaliella salina* CCAP 19/18. Healthy cultures *Nannochloropsis, Chlamydomonas*, and *Dunaliella* were suspended in 70% ethanol and frozen at -20°C for at least 24h. As UTEX 3031 often grows in tetrads, cultures were subjected to 10% N stress to induce single cell growth then frozen in 70% ethanol, following Sanders et. al. (Sanders, 2022). The cultures were then resuspended in BG11 growth media and transferred in triplicate to a 96-well plate with 199μL culture with 0.5μL Syto 9 (Invitrogen, Waltham, MA) and 1μL of 10mg/mL RNase (Thermo Scientific, Waltham, MA). The plate was incubated at 37°C for 30 minutes, then cooled for analysis measurement on a BD Accuri C6 Plus flow cytometer (BD Biosciences, San Jose, CA) equipped with 488 nm and 640 nm lasers. Fluorescence emission at 488 nm through the 533/30 filter was collected for 10,000 events at a 14 μL/sec flow rate. Standard protocol check to ensure instrument repeatability was done using analysis of Spherotech 8-Peak Validation Beads (BD Biosciences, San Jose, CA, #653144). Flow cytometry data was analyzed using BD CSampler Plus Analysis software to take the mean fluorescence of each peak of the fluorescence versus count histogram. Post-flow cytometry, samples were examined under a light microscope to check cell health and growth in single cells versus doublets or tetrads. Statistical analyses were conducted using R version 4.1.2 (R Core Team) and figures were produced using the package ggplot2 (Wickham, 2016, 2). A linear model was used to predict relative fluorescence units (RFU) based on the genome size in megabases (Mbp) from *Nannochloropsis* and *Chlamydomonas*. CCAP 19/18 was excluded from the model because the fluorescence was inexplicably high in multiple cultures.

### PacBio HiFi Library Preparation

Two libraries were sequenced on the Sequel and Sequel II respectively, as the same Sequel instrument was upgraded to a Sequel II during the project. The gDNA was sheared to the target size of 15 Kbp on Megaruptor 2 and purified with AMPure PB beads. HiFi SMRT libraries were prepared according to the Pacific Biosciences protocol. Single strand overhangs were removed, followed by the DNA damage repair, end repair and A-tailing reactions. Overhang adapters were ligated, followed by a nuclease treatment reaction to remove damaged DNA and misligated products. For sequencing on the Sequel instrument the SMRT bell libraries were purified and size selected using diluted AMPure PB beads. Sequencing primer v.4 was annealed and Sequel DNA polymerase 3.0 bound to the SMRT bells. Diluted SMRT bell/ DNA polymerase complex was sequenced on two SMRT cells 1M using the sequencing plate 3.0 and 20 hour movies.

### Hi-C Library Preparation

To build chromosome-level scaffolds, the “Proximo” service (https://phasegenomics.com/products/proximo/) from Phase Genomics (Seattle, WA) was utilized to generate Hi-C sequencing reads. A total volume of 5 mL of fresh, wet pelleted *S. obliquus* UTEX 3031 cells were provided to Phase Genomics for analysis. Cells provided were pelleted at 3000 × g for 5 minutes and supernatant was removed prior to shipment. The Proximo protocol generated approximately 172 million Illumina paired reads (2 × 75bp reads) from the Hi-C library generation and sequencing process.

### Oxford Nanopore Library Preparation

The concentration of gDNA was obtained using the Qubit DNA Assay Kit (ThermoFisher Scientific, Cat. # Q32854). The size of the DNA was determined using the Tapestation gDNA assay (Agilent, Cat. #5067-5366 and 5067-5365). A MinION library was prepared from 1 μg of DNA using the Ligation Sequencing Kit (Oxford Nanopore, Cat. #SQK-LSK109). The DNA was repaired using the NEBNext Companion Module for ONT Ligation Sequencing (NEB, Cat. #E7180S) and adapters were ligated onto the ends of the fragments to generate a library. The library was purified using AMPure XP beads (Beckman, Cat. #A63381), quantified using the Qubit 1x dsDNA HS Assay (ThermoFisher Scientific, Cat. #Q33231), and was sequenced on one R9.4.1 flow cell (Oxford Nanopore, Cat. #FLO-MIN106) for 48 hours.

### Genome Assembly

The Illumina reads from the Proximo HiC assay were trimmed and quality filtered (“-q 20”) using fastp v. 0.20.1 (Chen et al., 2018). The raw PacBio subreads were converted into Circular Consensus Sequences (CCS or “HiFi”) reads using PacBio’s CCS module (https://ccs.how.) This resulted in 1,864,935 reads, totaling 18,370,360,087 bp, or approximately 89x coverage per haplotype (assuming a diploid size of ∼206 Mbp.) The CCS reads were assembled using hifiasm v. 0.15.5 (Cheng et al., 2022) with parameters “--f-perturb 0.99 --n-weight 6 --n-perturb 1000000” and phased into individual haplotypes using hifiasm’s Hi-C phasing module. The two haplotypes were concatenated into a single file and any contigs less than 50 Kbp were removed. The remaining contigs were scaffolded using the same Hi-C data in conjunction with the Juicer (Durand et al., 2016) + 3d-dna pipeline (Dudchenko et al., 2017). Using Juicebox, assembly errors such as misjoins and false inversions were manually fixed per the manual instructions, and the assembly was scaffolded into individual chromosomes. The resulting .assembly file was exported and converted back into a .fasta file using juicebox_assembly_converter.py from Phase Genomics (https://github.com/phasegenomics/juicebox_scripts/blob/master/README.md.)

### Gap Closing and Quality Assessment

Four false duplications and five mis-assemblies were fixed by performing a local reassembly with hifiasm. To fill the 22 gaps caused by long tandem repeats (LTRs), we followed part of the approach by *Nurk et. al*., *Page 4 (Nurk et al., 2022)*. Specifically, HiFi graphs were generated to guide the correct walk/path using both CCS reads and ONT reads. First, with the CCS reads, we reconstructed a local de Bruijn graph using MBG (Rautiainen and Marschall, 2021). The k-mer size used to generate the graphs varied, but were usually k=1501, 2501, or 3501. Then, GraphAligner (Rautiainen and Marschall, 2020) was used with parameters “-x dbg” to align raw Oxford Nanopore Technologies (ONT) reads to the graph. GraphAligner generates a file which shows the sequence name and an alignment which shows the path the ONT read traverses along the graph. In addition, we used Bandage (Wick et al., 2015) to visualize the de Brujin graphs and copy sequences directly from the graph. Once the correct path was found, the path was pasted into Bandage, the sequence copied, and manually inserted into the .fasta file. Gaps caused by (TG_n_) dinucleotide repeats resolved using single HiFi reads and corrected Nanopore reads that spanned the gap.

Haplotype assembly completeness was assessed using BUSCO (Simão et al., 2015) version 4.1.4 with the Chlorophyta (ODB10) database, which includes 1,519 genes built from 16 Chlorophyte genomes. Additionally, genome-wide read coverage was analyzed. In brief, a 10 Kbp non-overlapping bin BED file was created by using “bedtools makewindows” (Quinlan and Hall, 2010) with the parameter “-w 10000”. Next, coverage for each bin was calculated using “bedtools coverage” with parameter “-mean” using the aforementioned BED file and the read alignments in BAM format. The resulting coverage information/maps were analyzed with plots created using custom R scripts and the read pileups in corrected regions were visually inspected in Tablet (Milne et al., 2010) to verify that coverage levels were consistent with neighboring unique sequences around the gap. Regions determined to be differential between the Illumina sequencing performed in 2017 and the PacBio sequencing performed in 2020 and 2021 were verified by polymerase chain reaction (PCR) using RedTaq (Sigma-Aldrich) according to standard protocols. For chromosome 13, a single forward primer upstream of the 1 Mbp region that is absent in chromosome 13B (GAACAGGGCCATCAACACAC) was used to coamplify the two possible DNA conformations with a reverse primer present in the region (CAAATGGGATGAGTGAAAACC) and after the deleted region (GCCTCCACAATCATCGTCTT). For chromosome 2, three independent regions were amplified to confirm the conformation of the assembled chromosome sequences.

Primer sets 1 (Chromosome 2A: 7215024-7217024, FP: GGGTACAGAAGCAGGGGTAA, RP: TCCATGTCAGACCACTCACA), 2 (Chromosome 2A: 8141992-8143992, FP: TTTCCTGCAAGCACCAACAA, RP: GACCAAGAGCAGCATCAAGG), and 3 (Chromosome 2B 7053654-7055654, FP: ACATGTTGCGTTCCTTCCTG, RP: GATTGGCAGGCGTGATGAG) successfully amplified the regions of interest. All primer sets were confirmed to uniquely target a single genomic sequence and all reported bands match the theoretical size of the region of interest (Supplemental Figure S7).

### Structural Annotation

Structural gene models were constructed using a modified version of BRAKER v2.1.6 (Bruna et al., 2021). Specifically, we trained repeat models of the genome assembly using RepeatModeler v 2.0.2 (Flynn et al., 2020) and masked repetitive regions using RepeatMasker v4.1.2 (Chen, 2004). Paired-end, poly-A selected RNA sequencing was performed using an Illumina NextSeq 500. Individual libraries consisted of ambient air and CO_2_ supplemented grown samples at log and mid-log growth intervals for a total of 123,614,170 paired-end 150bp RNA-seq reads encompassing a diverse transcript pool (Jenna Y Schambach et al., 2022). In parallel, we combined raw RNA-seq reads from all libraries and trimmed them using fastp v0.23.2 (Chen et al., 2018). Trimmed reads were aligned to the genome assembly using HISAT2 v2.2.1 (Kim et al., 2019, 2), and subsequently sorted using Samtools v1.14 (Li et al., 2009). Using the soft-masked genome assembly and the transcriptomic evidence, we performed structural annotation using BRAKER v2.1.6 (Bruna et al., 2021) with “--softmasking” option. As a part of this pipeline, GeneMark-ES and AUGUSTUS versions corresponding to the BRAKER v2.1.6 were integrated to process the transcriptomic evidence and to utilize it to train a species-specific gene model (Lomsadze et al., 2005; Stanke et al., 2006). In addition, we predicted non-coding RNAs in nuclear and organelle genomes. tRNAs were predicted using tRNAscan-SE v2.0.9 (Chan et al., 2021) with default parameters. rRNAs were predicted using barrnap v0.9 (Seemann, 2022) with default parameters.

### Functional Annotation

The putative functions of the gene models were annotated using homology searched against the CDD, Pfam, PIRSF, TIGRFAM, and SWISSPROT databases (Accessed at January 2022) **(Barker et al., 2000; Boeckmann et al., 2003; Haft et al., 2003; El-Gebali et al., 2019; Lu et al., 2020)**. Interproscan v5.46-81.0 (Jones et al., 2014) was utilized to perform annotation of protein functions and linked Gene Ontology terms (Gene Ontology Consortium, 2015) against CDD, Pfam, PIRSF, and TIGRFAM databases. BLASTP v2.10.0 (Camacho et al., 2009) was used to search protein queries against SWISSPROT databases with E-value cutoff of 1e-6 and enablement of SEG filtering (Wootton and Federhen, 1996) and soft-masked region filtering. Only the top BLAST hit per each protein query was selected. The KEGG Automatic Annotation Server (KAAS) (Moriya et al., 2007) was used to associate structural-annotation-identified genes with KEGG Orthology (KO) identifiers (Kanehisa, 2019; Kanehisa et al., 2021). The KEGG Brite Database was then utilized to annotate KO ids with brite hierarchy, KO description, EC id, and gene symbols. Finally, overlapping genes were trimmed using ‘agat_convert_sp_gxf2gxf.pl’ and ‘agat_sp_fix_overlaping_genes.pl’ scripts of AGAT package v0.8.0 (Dainat et al., 2022).

### Synteny and Genomic Rearrangement Analysis

FastANI (Jain et al., 2018) with default parameters was used to calculate the average nucleotide identity (“ANI”) between each pair of chromosomes. For each pair, the A haplotype was used as the reference and the B haplotype was used as the query. Synteny between the UTEX 3031 haplotypes and between other *Scenedesmus* strains was measured first by completing a whole-genome alignment with nucmer version 3.1 (Kurtz et al., 2004). Next, SyRI v1.6 (Goel et al., 2019) was utilized with the NUCmer output to detect large-scaled genomic rearrangements (GR) and syntenic regions. All detected rearrangements and syntenic regions were plotted using plotsr v0.5.3 (Goel and Schneeberger, 2022).

After syntenic regions and unaligned regions were estimated, we performed downstream GR quantification between the ‘A’ haplotypes as the ‘reference’ and all other haplotypes of chromosomes as ‘query’. GRs smaller than 500 bp were excluded. Genes overlapping with GRs were also summarized. Here, we identified clusters of multiple GRs that are located nearby (<1 Kbp) in both reference and query sequences, referred to as GR blocks. We selected and visualized long GR blocks with a size of at least 100 Kbp.

Deleted or translocated genes were verified for each haplotype by conducting homology searches across the entire genome assembly. A gene was categorized as ‘unique’ using the following criteria: tblastn (qcovs >= 50, length >= 40, evalue <= 1e-5) and reciprocal best hit (length >= 40, evalue <= 1e-10) comparisons against haplotype assemblies (Camacho et al., 2009). Translocations were identified as reciprocal best hits in all-vs-all comparisons via minimap2 (Li, 2018, 2) (default parameters, input sequences were intron inclusive gene coding regions). Functional pathway enrichment of GO terms was performed using the R package TopGO (Alexa and Rahnenfuhrer) function ‘runTest’ (algorithm=“weight01”, statistic=“fisher”).

### Intron Analysis

Introns were extracted from the .gff3 annotation of UTEX 3031 for all haplotypes. blastn (Altschul et al., 1990) was used with an E-value cutoff of 1*10^−30^ to search/compare introns identified in the genome assemblies of four *Scenedesmus* strains (DOE0013, EN004, UTEX1450, UTEX2630). Custom perl and R scripts were used to identify which strain of *Scenedesmus* each intron in UTEX 3031 assembly was most closely related to, based on e-values. For this process, e-values of 0.0 were converted to 1*10^−180^ and exact equal e-values between an intron from UTEX 3031 and any two strains were removed from the analysis.

### Repeat identification and mapping

First, kmc v 3.1.1 (Kokot et al., 2017) was used with parameters “-fa, -k151, -cs100000” to count k-mers in the genome. Next, the resulting .kmc file was converted into a .txt file using kmc_dump with default parameters. Then, Satellite Repeat Finder (aka “srf”) (https://github.com/lh3/srf) was used to generate a FASTA file with the identified repeats. Repeats less than 50 bp were filtered out of the FASTA file. Finally, the remaining repeats were aligned to the entire genome using minimap2.

### CpG Methylation Calling and Analysis

Kinetics information was extracted from the PacBio HiFi reads using the “--hifi-kinetics” parameter in PacBio’s ccs tool (https://ccs.how). The kinetics information was then fed into the PacBio tool “primrose” (v1.3.0) to predict 5mC methylated sites on each CpG motif. Next, the BAM output from primrose was analyzed using “aligned_bam_to_cpg_scores.py” script from PacBio’s repository (https://github.com/PacificBiosciences/pb-CpG-tools), which outputs a BED file containing CpG positions and their respective probabilities of 5mC methylation.

To identify putative promoter regions, a heuristic was created to classify genome regions with different methylation probabilities. First, we identified positions in the genome which had a methylation probability of > 60%. Next, for each position, the distance was calculated between the identified position and the next methylated position. If this distance was greater than 350 bp, the region was marked as a putative promoter. For each gene in the genome annotation, the distance was measured between the start of transcription and the beginning of each identified promoter region. If the distance from the transcription start site to the nearest putative promoter was less than 2000bp, the gene was considered to contain a hypomethylated and transcriptionally active promoter region. Genes lacking a hypomethylated promoter were considered to be transcriptionally inactive.

## Accession numbers

Raw reads are available in the sequence read archive (SRA) under PRJNA826445 with the SRA numbers SRR21831894-SRR21831912. Full gene annotations as well as haplotype-specific and full genome assemblies are available on Phycocosm (link to be added upon acceptance). Each haplotype assembly of the complete genome is available at the NCBI GenBank database is deposited under PRJNA887759 (haplotype A) and PRJNA887758 (haplotype B).

## Funding

This work was funded by the U.S. Department of Energy Office of Energy Efficiency and Renewable Energy, Bioenergy Technologies Office to S.R.S. (contract NL0029949, WBS 1.3.1.600).

## Acknowledgements and Author Contributions

Special Thanks to the LANL Genomics and Bioanalytics Group for valuable feedback. This work has been cleared for public release by Los Alamos National Laboratory (LA-UR-XX-XXXX). T.C.B. completed the genome assembly, including scaffolding and gap closing; calculated genome assembly statistics; produced synteny and genomic rearrangement data; calculated coverage and ANI for each chromosome; produced 5mC methylation data; created heuristic for detecting hypomethylated areas. E.R.H. performed genomic analyses and cellular DNA quantitation. T.K. and W.E. performed structural and functional annotation of the genome assembly. T.K. performed and interpreted genomic rearrangement analyses. C.K. co-developed and interpreted enrichment and functional annotation analyses, secured RNA-seq data, designed and performed PCR validation of large chromosomal differences between pairs, aided in interpretation of coverage anomalies, and contributed to algorithm development, analysis, and interpretation of methylation data. S.I.K. identified unique genes, co-developed functional annotation analysis, developed enrichment analyses of unique genes, translocations, deletions, methylated regions, and genomic rearrangement blocks, and identified translocations. Y.K. performed the PacBio library preparations and sequencing data. C.D.G. prepared genomic DNA libraries and generated the MinION sequencing data. K.T.Y.M. performed cellular DNA quantitation. J.P contributed to experimental planning, comparative genome analysis, and data analysis. B.T.H. contributed to the experimental planning, genome assembly, and annotation. S.R.S. designed, conceived, and supervised this study. All authors contributed to the writing and editing of this article. The authors declare no potential financial or other interests that could be perceived to influence the outcomes of the research.

## Ethics declarations

### Ethics approval and consent to participate

Not applicable.

### Consent for publication

Not applicable.

### Competing interests

The authors declare that they have no competing interests.

---

**Supplemental Figure S2.**
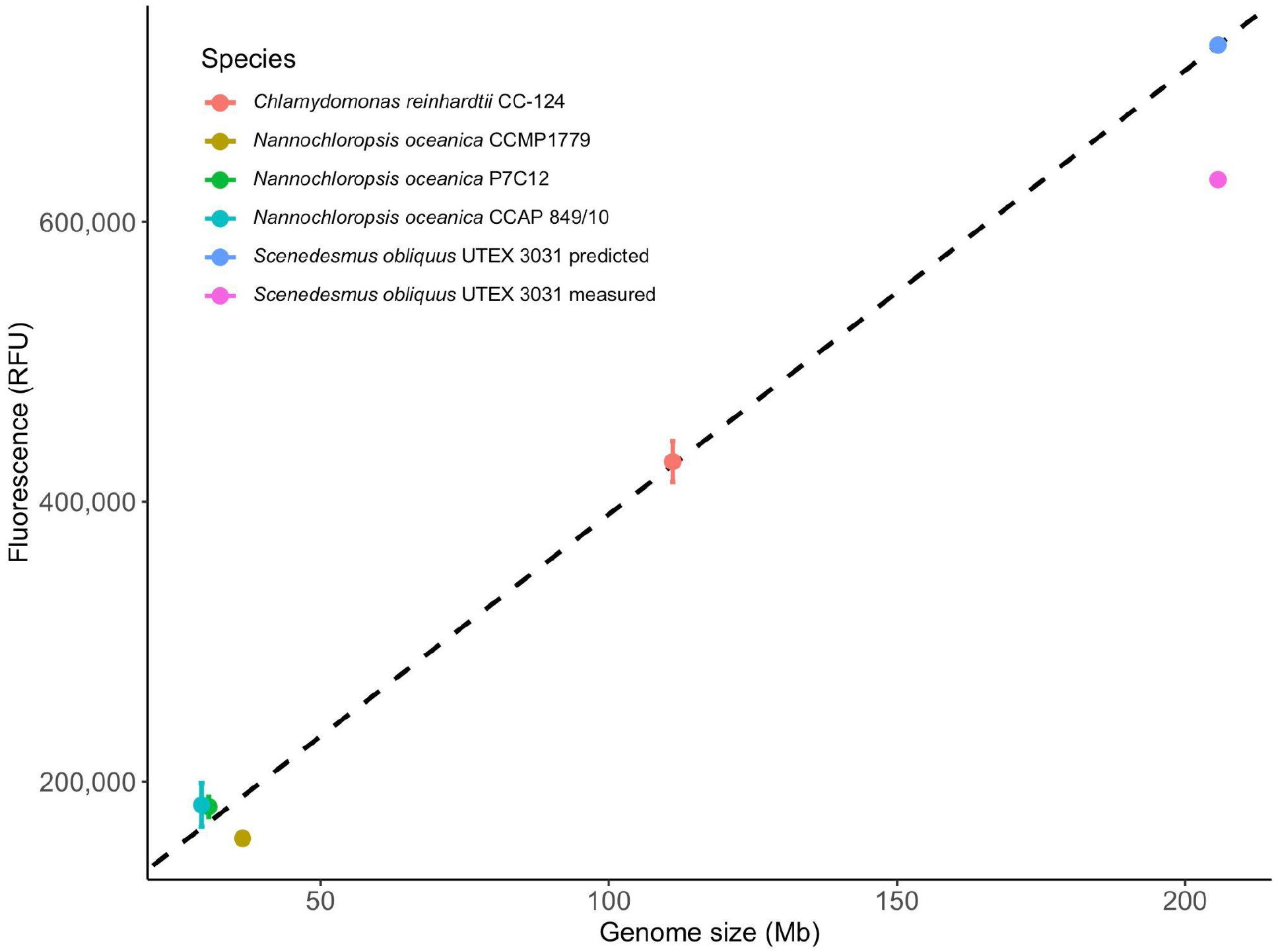
Relative fluorescence units (RFU, mean± standard error) of Syto 9 stained cells of 6 algal species and strains and their genome sizes (Mb) with a linear regression.

**Supplemental Figure S3.**
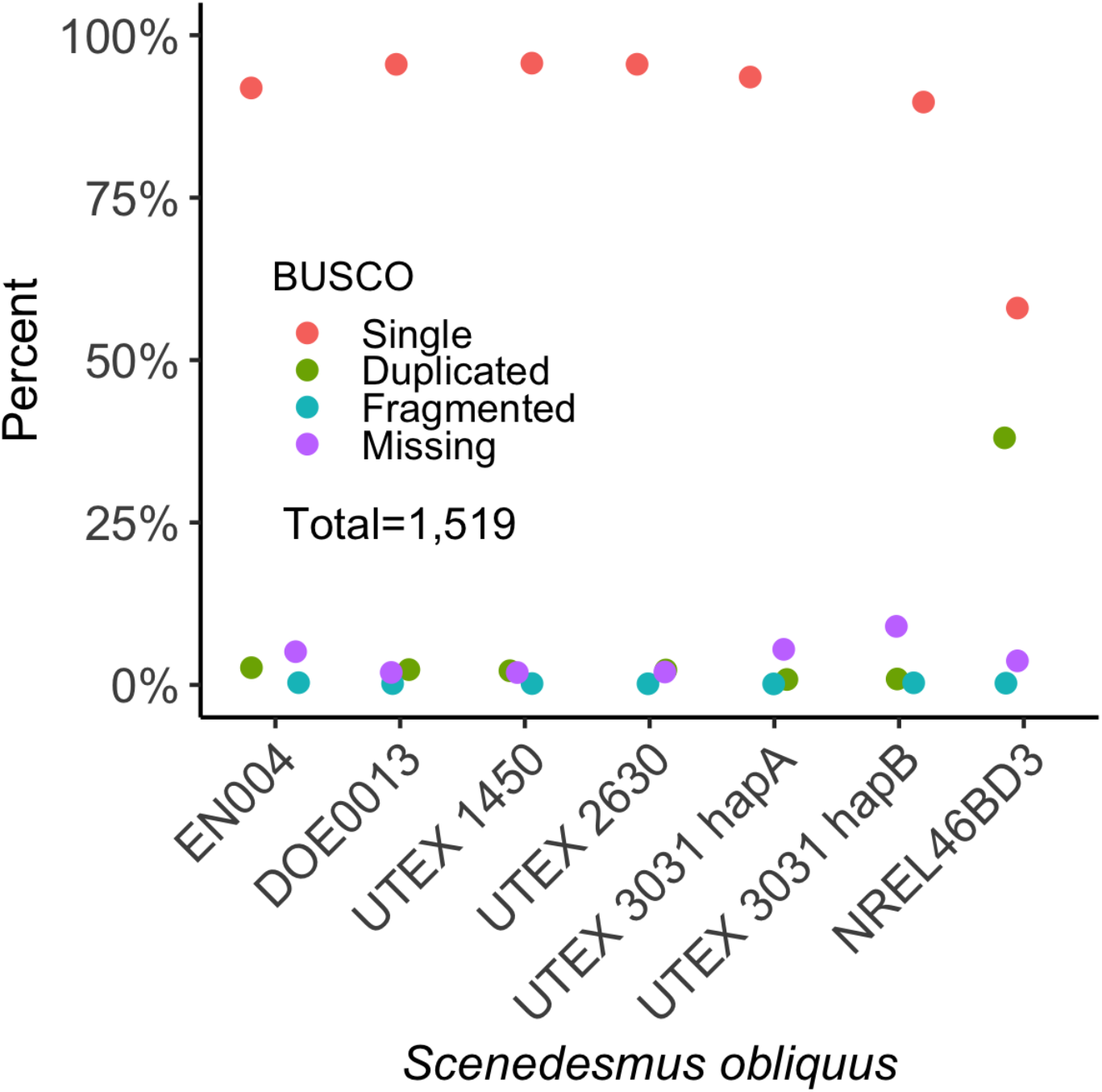
Presence of BUSCO (Benchmarking Universal Single-Copy Orthologs) in *Scenedesmus obliquus* genomes using the Chlorophyta gene set (n=1,519). Both haplotypes A and B have near complete representation of single copy orthologs.

**Supplemental Figure S4.**
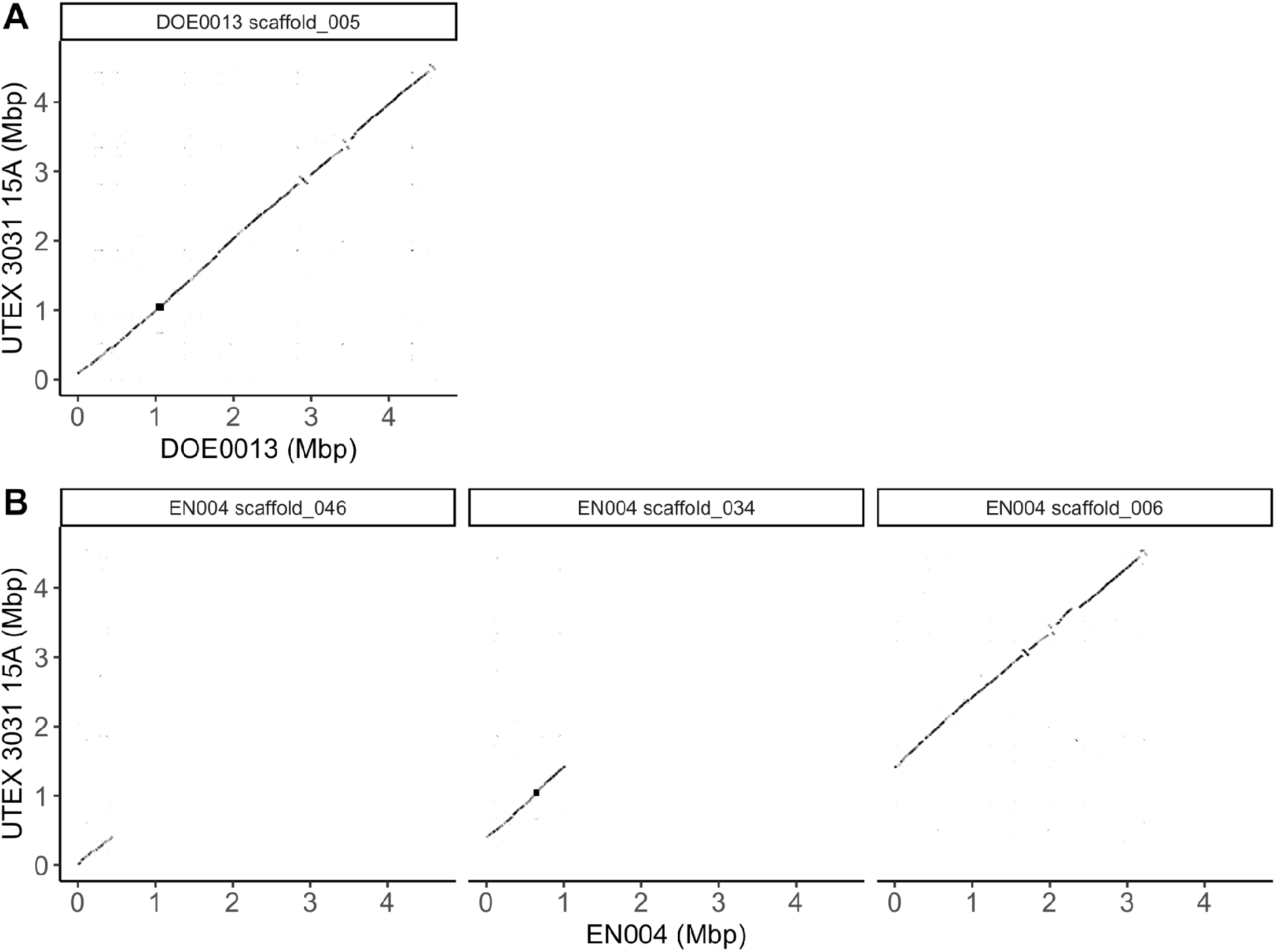
Homology between chromosome 15 (haplotype A) and scaffolds from DOE0013 and EN004 assemblies. A. DOE0013 scaffold 5 is nearly complete. B. Chromosome 15 is assembled into three scaffolds (46, 34, and 6) in EN004. The large square region near 1 Mb on UTEX 3031 chromosome 15a is a centromere. Three rearrangement differences between DOE0013 and UTEX 3031 (2.8 - 4.2 Mbp) are shared between EN004 and DOE0013.

**Supplemental Figure S5.**
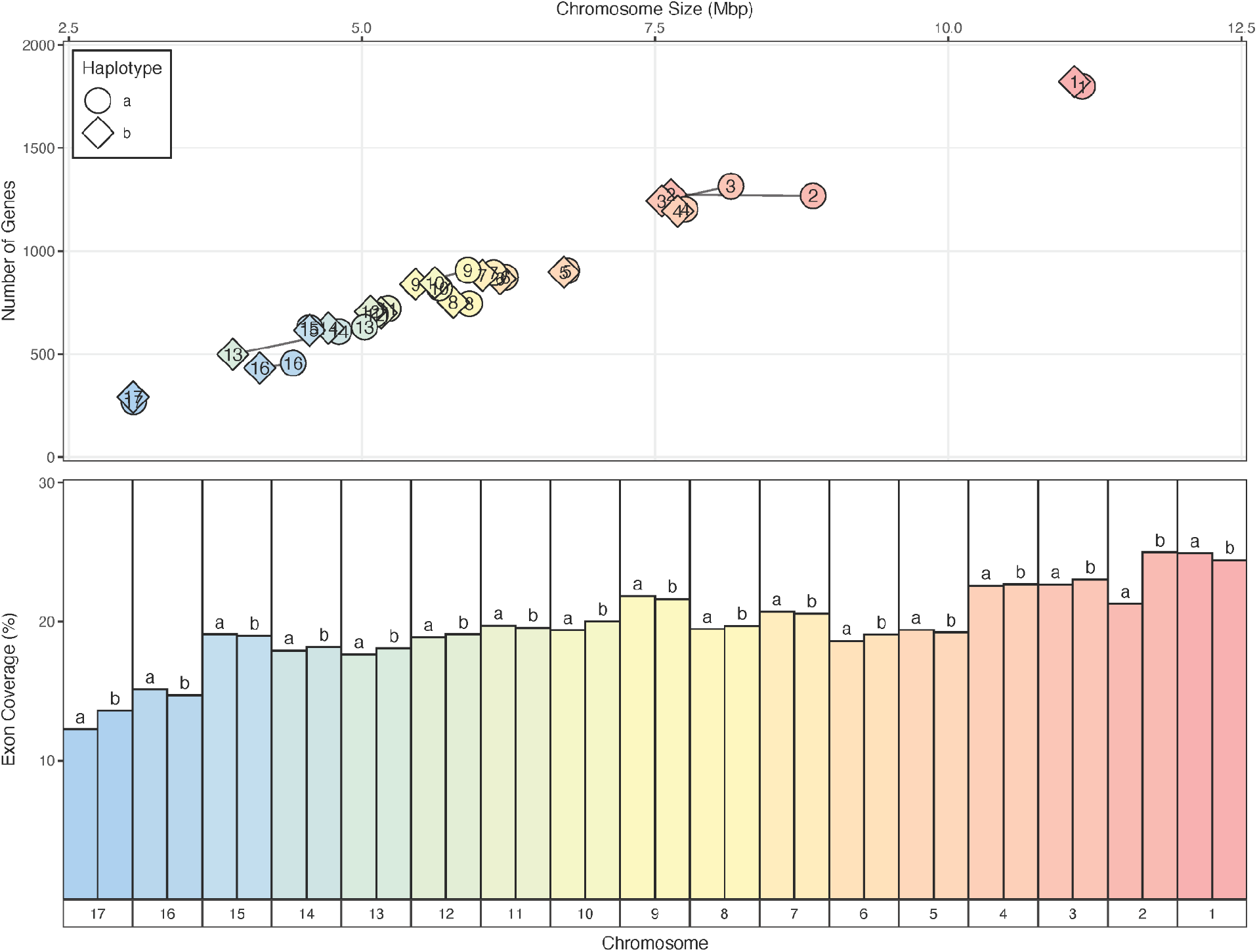
Chromosome Gene Count and Gene Density. The annotated gene count is plotted against total chromosome size for each chromosome pair (top). Lines connect pairs of haplotypes. Average gene length (bottom) was also calculated for each chromosome to identify potential gene length biases resulting in increased coding gene contribution.

**Supplemental Figure S6.**
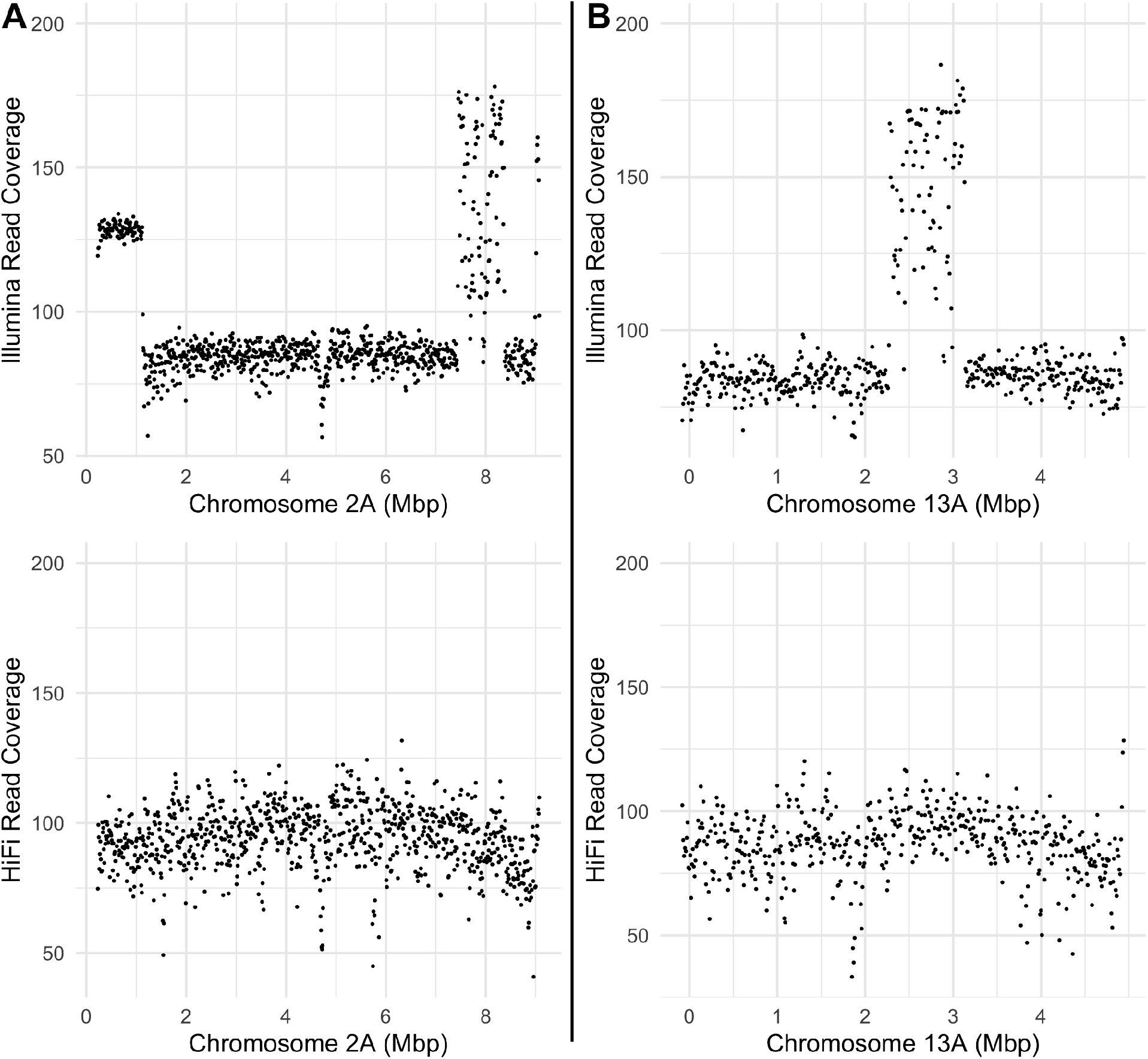
Read Coverage Analysis of Recent Deletions in Two Chromosomes. A) Illumina read coverage and PacBio HiFi read coverage of Chromosome 2A (A) and 13A (B), respectively after gap closing.

**Supplemental Figure S7.**
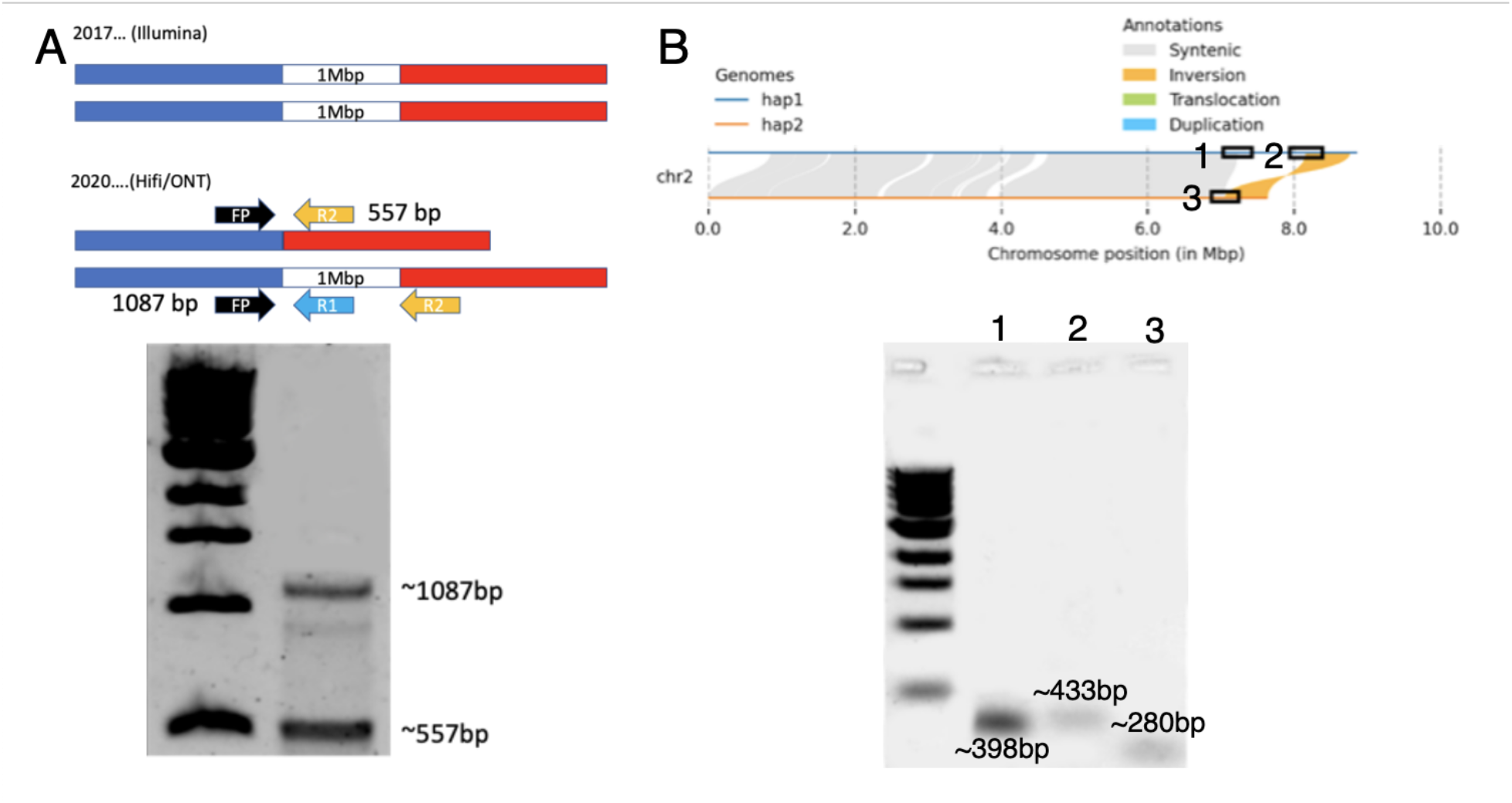
PCR design and gel size estimation of amplified products validated the deletion which occurred in chromosome 13B as well as the deletion and rearrangement of chromosome 2A and 2B.

**Supplemental Figure S8.**
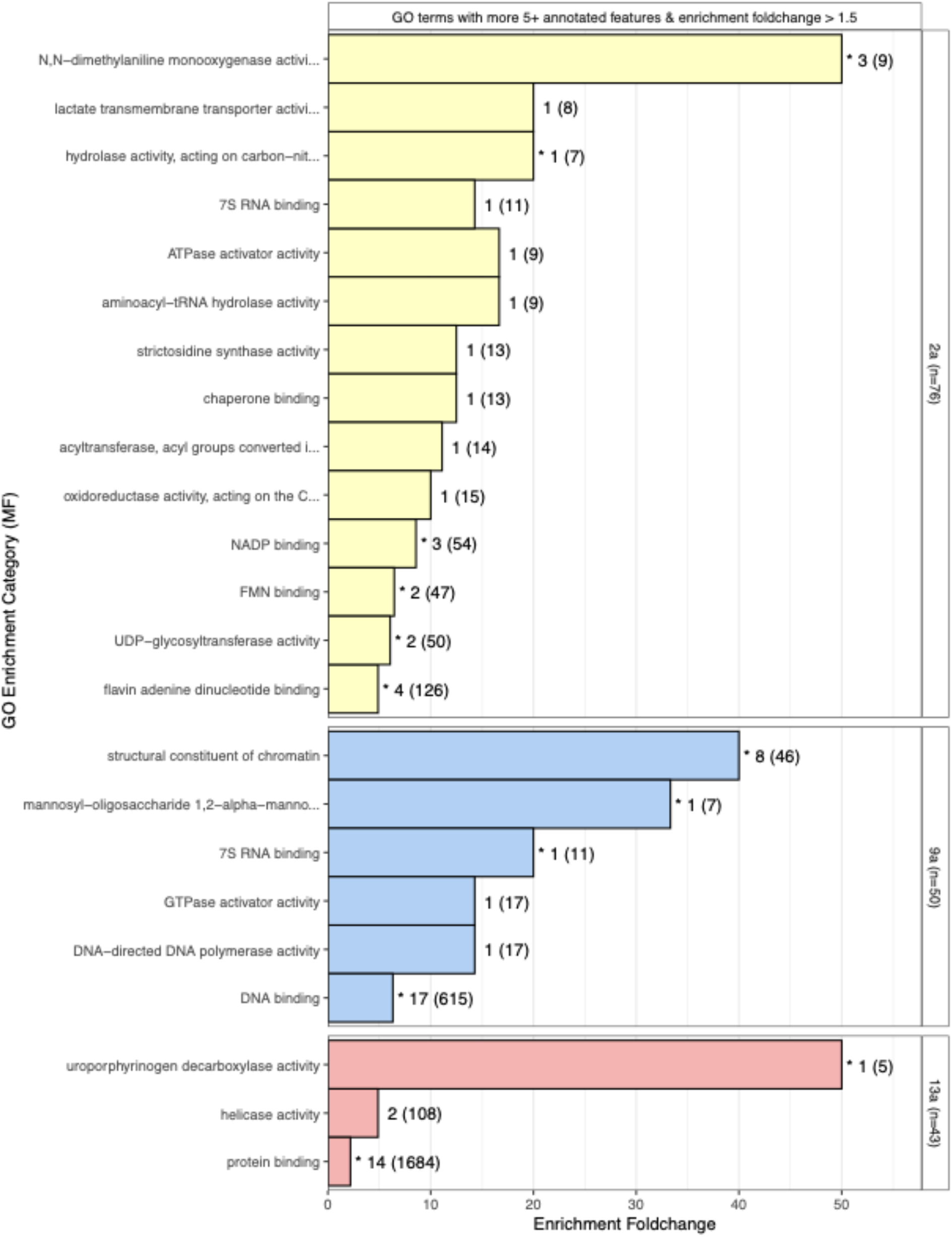
Enriched molecular functions across the three large chromosome pair indels (chromosome 2, 9, 13) with individual labels identifying the set size of genes in a given term on the indel region and the full term set size in the genome in parenthesis for reference. * = significant by fisher exact test (P≤ 0.05). Only terms with atleast 5 genes in the total GO term and with at least 1.5 fold increased abundance over statistically expected set size were plotted.

**Supplemental Figure S9.**
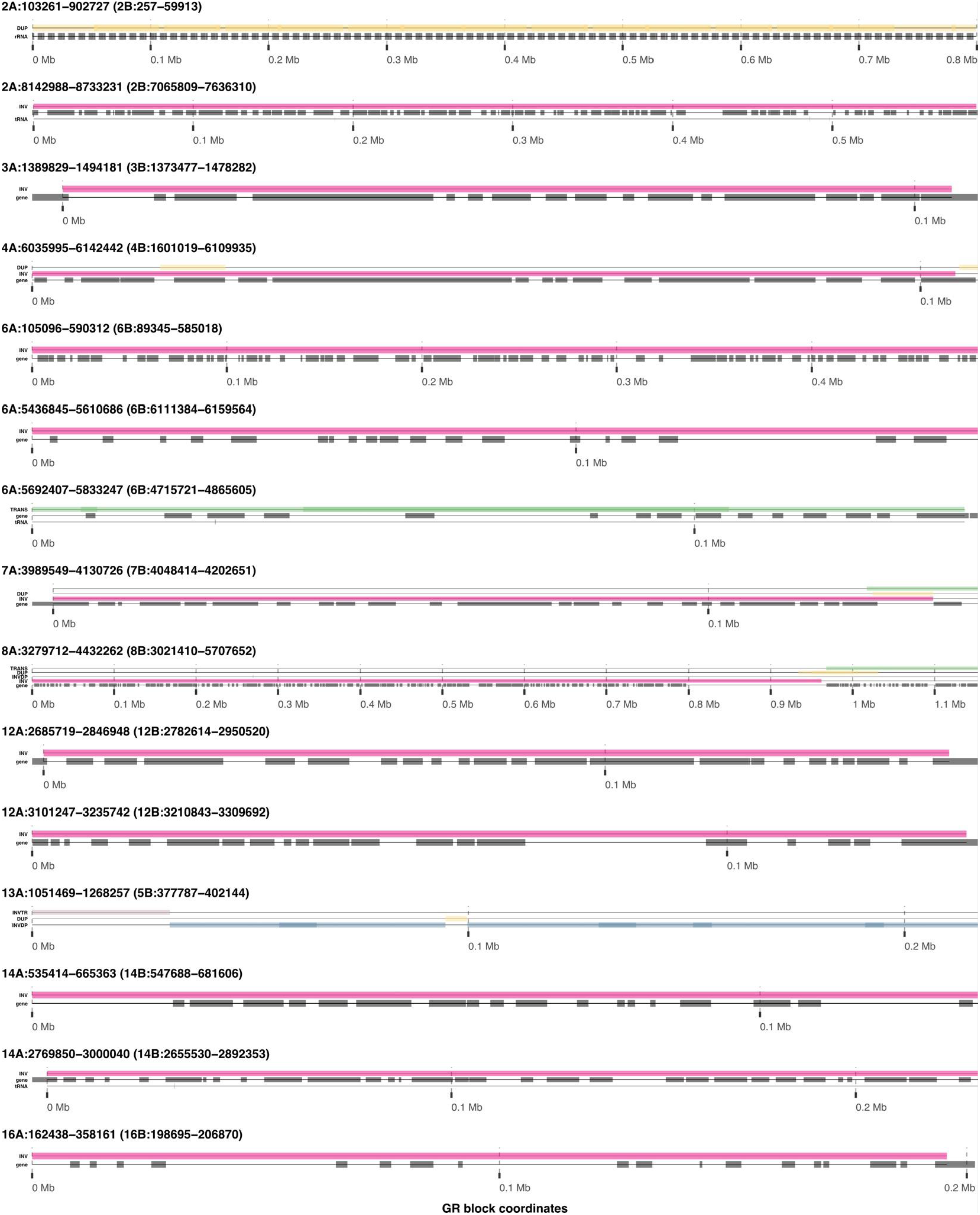
Overview of 15 long genomic rearrangement blocks. Different classes of genomic rearrangements are illustrated with colored rectangles (type labels at left of each rectangle), while protein coding genes, rRNA genes, and tRNA genes are illustrated with black rectangles.

**Supplemental Figure S10.**
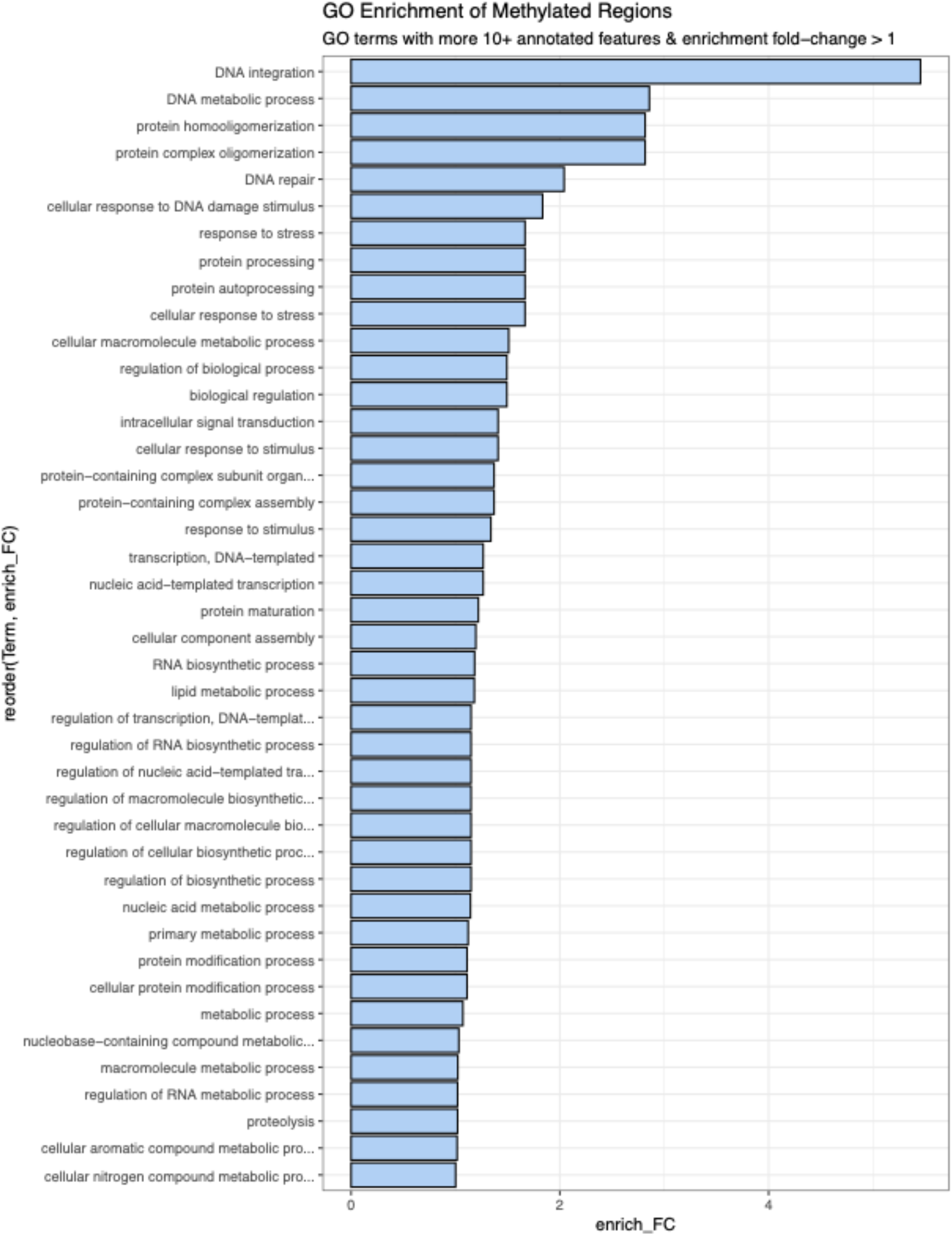
Enrichment of biological processes within the genes identified to contain hypermethylated promoter regions based on CpG methylation data.

**Supplemental Figure S11.**
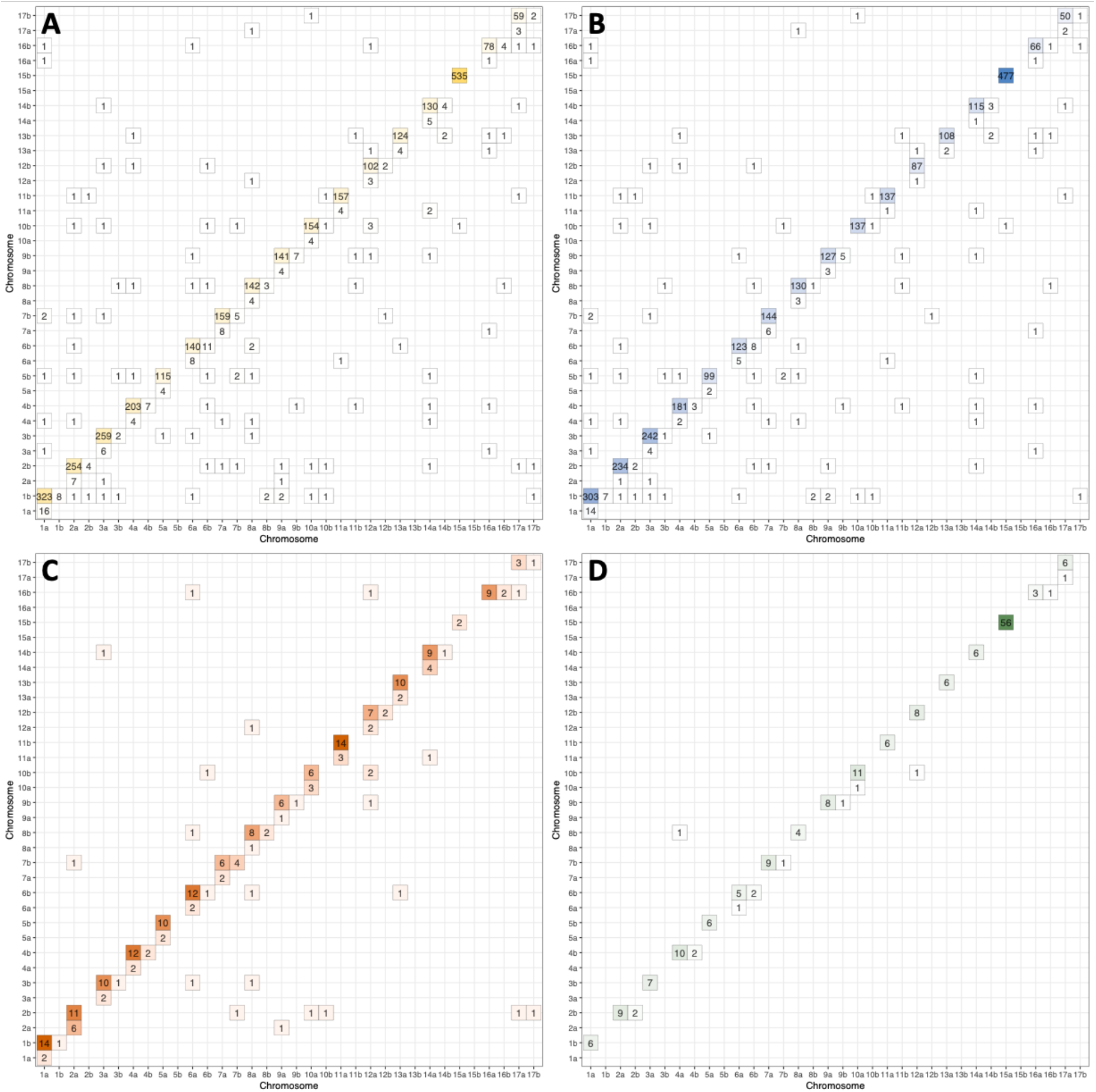
A) Distribution of all gene pairs, defined as reciprocal best hits with >90% sequence similarity. B) Distribution of non-methylated gene pairs. C) Distribution of 1/2-methylated gene pairs. D) Distribution of both-methylated gene pairs.

**Supplemental Figure S12.**
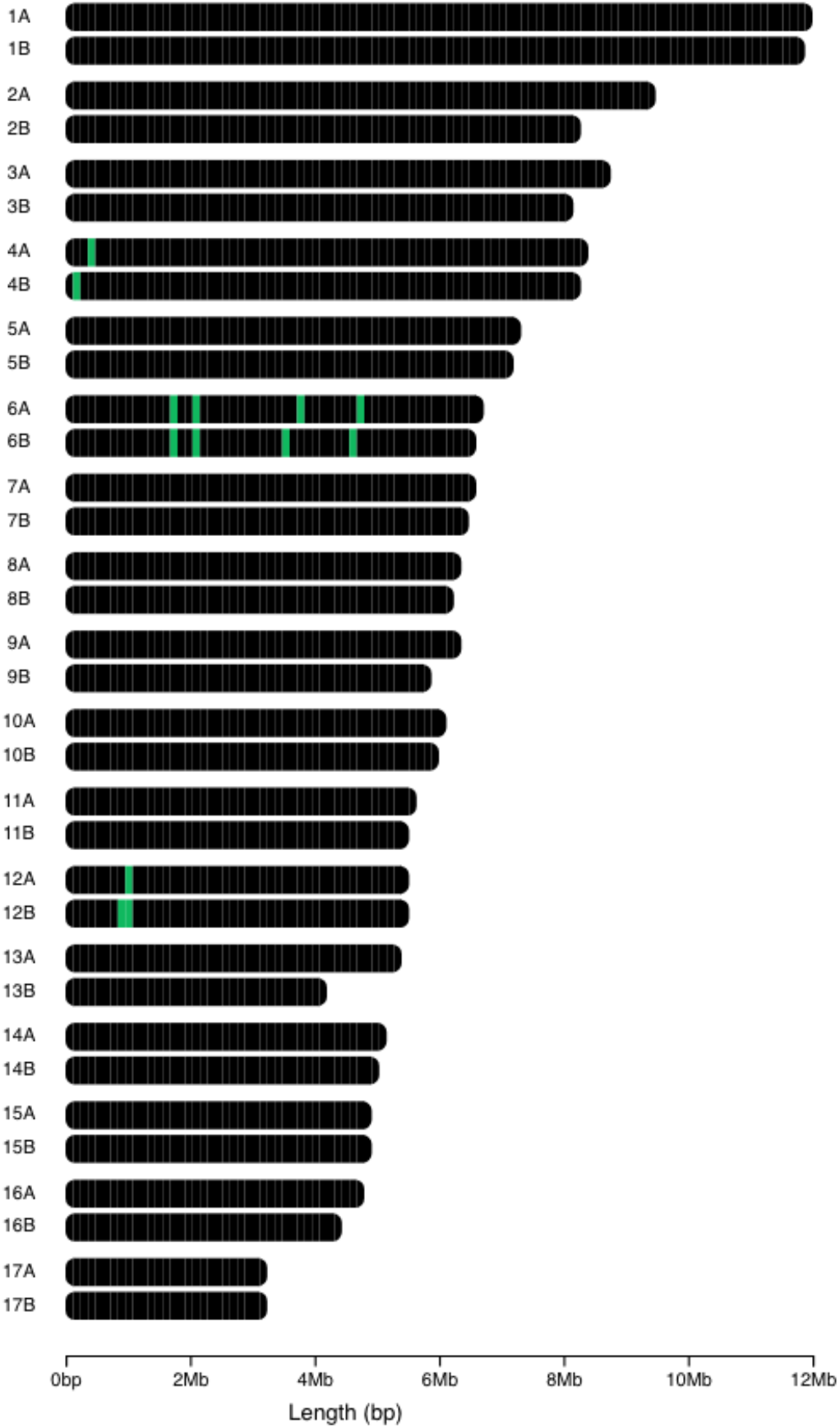
Locations of all genes (green) with RWP-RK functional domains in the genome of *Scenedesmus obliquus* UTEX 3031.

**Supplemental Figure S13.**
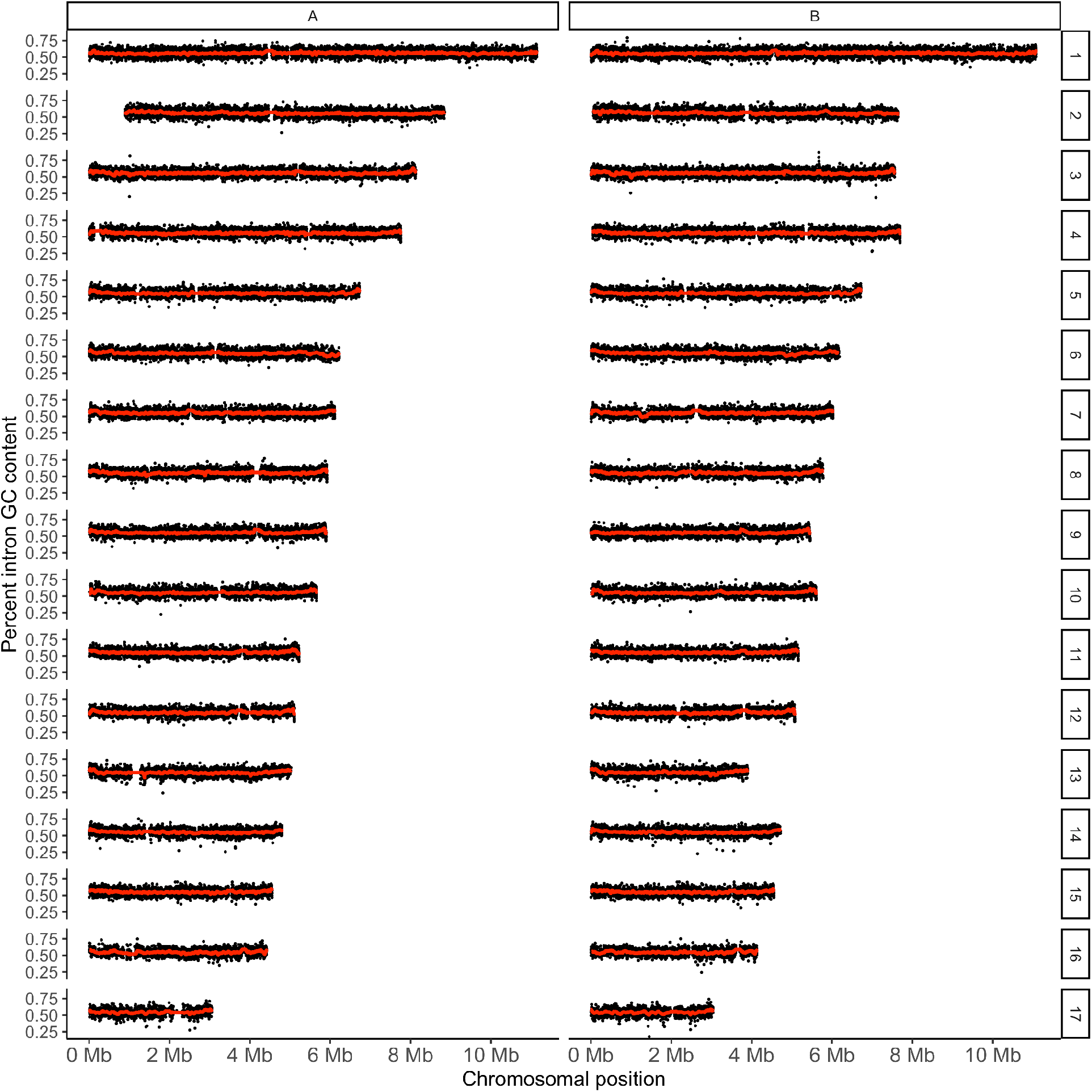
GC content of introns across all chromosomes. The red line indicates a rolling average of 50 introns.

## Notes

### Competing Interest Statement

The authors have declared no competing interest.

